# Polycomb repression works without Siesta

**DOI:** 10.1101/2025.07.18.664654

**Authors:** Tatyana G. Kahn, Andres Garrido, Anastasiya Yushkova, Maria Kim, Alexander Glotov, Sweda Sreekumar, Jan Larsson, Yuri B. Schwartz

## Abstract

Polycomb group proteins mediate epigenetic repression via multi-subunit complexes, including canonical Polycomb Repressive Complex 1 (PRC1), which monoubiquitylates histone H2A and binds histone H3 tri-methylated at lysine 27 (H3K27me3). The RING1 subunit of PRC1, critical for H2A ubiquitylation, forms other complexes. These variant RING1 complexes also ubiquitylate H2A but cannot bind H3K27me3, and their evolutionary origins and functions are debated. Using *Drosophila* genetics, we discovered that canonical PRC1 and variant RING1 complexes ubiquitylate H2A at distinct genomic regions. We established that the sole *Drosophila* PCGF protein specific for variant RING1 complexes, which we named Siesta, is not required for epigenetic repression of developmental genes, but controls larval locomotion independently of H2A ubiquitylation. Leveraging a massively parallel transgenic approach, we demonstrated that H2A ubiquitylation has minimal impact on transcriptional repression. Our findings imply that variant RING1 complexes operate outside the Polycomb regulatory system and that the popular PRC1 nomenclature needs revision.

## INTRODUCTION

Polycomb group proteins are well-known for epigenetic repression of specific master-regulatory genes, thereby ensuring that cells maintain their developmental identities. These proteins act as multi-subunit complexes, traditionally classified into two families: Polycomb Repressive Complex 1 (PRC1) and Polycomb Repressive Complex 2 (PRC2). The originally characterised PRC1 complex (purified from *Drosophila melanogaster* cells and sometimes referred to as canonical) consisted of Polycomb (Pc), Pleiohomeotic (Ph), Sex combs on midleg (Scm) proteins, as well as the heterodimer between RING1 (the product of *Sex combs extra* (*Sce*) gene) and Posterior sex combs (Psc) proteins (Shao et al., 1999). The Pc subunit of the canonical PRC1 specifically binds to histone H3 tri-methylated at lysine 27 (H3K27me3) (Fischle et al., 2003), a modification produced by PRC2 complexes (Czermin et al., 2002; Muller et al., 2002). Genes repressed by the Polycomb system are embedded in broad chromatin domains enriched in H3K27me3 (Kahn et al., 2006; Schwartz et al., 2006). This histone modification acts as a molecular mark that helps to propagate the repressed state following DNA replication (Coleman and Struhl, 2017; Laprell et al., 2017).

Subsequent proteome analyses of RING1 partners in mouse and human cells (mammals have two closely related orthologs RING1 and RING2 encoded by the *RING1* and *RNF2* genes) revealed complexes analogous to canonical PRC1, but also a series of complexes lacking orthologues of Pc, Ph and Psc (Gao et al., 2012; Hauri et al., 2016; Kloet et al., 2016). These variant complexes centre around the heterodimer between RING1/2 orthologues and one of the four PCGF proteins (PCGF1, PCGF3, PCGF5, or PCGF6). These RING1/2 complexes incorporate RYBP (or a closely related YAF2 protein) instead of Pc orthologues. They also include additional subunits absent from canonical PRC1 that vary depending on the identity of the PCGF subunit.

Initially, variant RING1/2 complexes, referred to as non-canonical PRC1, were thought to have evolved from ancestral canonical PRC1, representing a diversification of the Polycomb system that facilitated the emergence of vertebrate-specific traits (Blackledge and Klose, 2021). However, systematic tracing of genes encoding core components of RING1/2 complexes in the genomes from diverse animal clades has shown that the RING1/2 complexes found in vertebrates appeared early in animal evolution, and some of them were subsequently lost in many lineages (Gahan et al., 2020).

Furthermore, the variant RING1/2-PCGF3 complex appears to be the ancestral complex from which all other variants, including canonical PRC1, have evolved (Gahan *et al*., 2020). Boosting this view, *Drosophila* analogues of all variant mammalian RING1/2 complexes were recently characterised biochemically (Kang et al., 2022).

The genes encoding RING1 and PCGF3 orthologues are present in species from *Filasterea* and *Choanoflagellata* clades, the closest unicellular relatives of animals (de Potter et al., 2023; Gahan *et al*., 2020). This argues that the ancestral RING1 and PCGF3 had functions distinct from epigenetic repression of developmental genes and the control of cell differentiation. The genetic studies in *Drosophila* and mice leave no doubt that the canonical PRC1, which first emerged in the metazoan lineage (de Potter *et al*., 2023; Gahan *et al*., 2020), is critical for the repression of developmental genes (Akasaka et al., 2001; Bel et al., 1998; Isono et al., 2005; Jürgens, 1985; Oktaba et al., 2008; van der Lugt et al., 1994). To what extent the variant metazoan RING1/2 complexes operate as part of the Polycomb system repressing developmental genes or, like the ancestral complexes, have unrelated functions, is less clear.

Functional studies provide mixed insights. In the developing mouse neocortex, genetic disruption of canonical PRC1 impairs the maintenance of cell lineage identity during neurogenesis and gliogenesis, whereas variant RING1/2 complexes appear to play a minor role (Hoffmann et al., 2025). Conversely, studies in mouse embryonic stem cells presented conflicting evidence. Some suggested that variant RING1/2 complexes synergise and are functionally more important than canonical PRC1 to repress developmental genes (Fursova et al., 2019). Others reported that canonical PRC1 plays a central role (Scelfo et al., 2019). Regardless, multiple lines of evidence suggest that at least some of the variant RING1/2 complexes have functions unrelated to epigenetic repression. For example, genetic ablation of RING1 and RING2 in mouse epidermal progenitor cells leads to skin fragility, which does not happen after removal of PRC2 (Cohen et al., 2019). Furthermore, in mouse brain cells, RING2-PCGF3 complexes bind in the vicinity of Transcription Start Sites (TSS) of a subset of active genes, where they appear to promote rather than repress transcription (Gao et al., 2014; Liu et al., 2021).

To what extent the catalytic activity of RING1 complexes is important for the repression of developmental genes is another open question. Both canonical PRC1 and variant RING1/RING2 complexes function as E3 ligases, monoubiquitylating histone H2A at a conserved residue, lysine 119 in mammals (H2AK119ub) and lysine 118 in *Drosophila* (H2AK118ub) (Gao *et al*., 2012; Tavares et al., 2012; Wang et al., 2004). Variant RING1/RING2 complexes seem to be more active E3 ligases *in vitro* compared to canonical PRC1 (Rose et al., 2016). This may be due to the differences in the biochemical properties of distinct PCGF proteins (Teslenko and Fierz, 2025) or additional stimulation by the RYBP/YAF2 subunit, which is unique to the variant complexes (Gao *et al*., 2012; Rose *et al*., 2016). With a size of ∼8.5 kDa, ubiquitin may physically affect chromatin structure. Thus, single-molecule magnetic tweezers experiments suggest that H2AK119ub stabilises *in vitro* reconstituted nucleosome particles by preventing the DNA unwrapping from the histone core (Xiao et al., 2020).

This may inhibit transcription. On the other hand, fluorescence resonance energy transfer measurements suggest that H2AK119ub interferes with chromatin fibre folding (Bonnet et al., 2022) making DNA more accessible to transcription regulators. In addition, H2AK119ub increases the affinity of PRC2 complexes containing accessory subunits Aebep2 and Jarid2, which, in turn, stimulates H3K27 methylation by this PRC2 variant (Kalb et al., 2014; Kasinath et al., 2021).

Consistent with its potential involvement in repression, H2AK119ub (and H2AK118ub in flies) is enriched within developmental genes repressed by the Polycomb system (Blackledge et al., 2020; Bonnet *et al*., 2022; Fursova *et al*., 2019; Kahn et al., 2016; Lee et al., 2015; Tamburri et al., 2020). In mouse embryonic stem cells, the replacement of RING1 and RING2 with a variant lacking E3 ligase activity led to a loss of H2AK119ub and a correlating increase in transcription of many developmental genes normally repressed by the Polycomb system (Blackledge *et al*., 2020; Tamburri *et al*., 2020).

However, the mechanistic link between the loss of H2AK119ub and de-repression remains ambiguous, as these cells also exhibited a loss of canonical PRC1, PRC2 and H3K27me3 from the same genes, making it difficult to isolate the specific contribution of H2A ubiquitylation. Remarkably, *Drosophila* mutants lacking RING1 E3 ligase activity or having histone H2A replaced with the truncated version that cannot be ubiquitylated show no obvious defects in repression by the Polycomb system (Pengelly et al., 2015). This argues that, even if H2AK119/K118 ubiquitylation contributes to repression in some species or cell types, it is not a universal mechanism by which the Polycomb system represses transcription of developmental genes.

Comparing how the epigenetic repression of developmental genes is affected by genetic ablation of variant RING1/RING2 complexes or canonical PRC1 is key to understanding their relative contributions. Removing either PCGF proteins specific for variant RING1/RING2 complexes or the PCGFs found in canonical PRC1 would be one way to accomplish this. Such a study is notoriously difficult to conduct in mammalian species because they have four PCGF proteins specific for variant RING1/RING2 complexes and two other PCGF proteins specific for PRC1, all encoded by separate genes located in different parts of the genome. The *Drosophila* model offers a powerful alternative to overcome this complexity. Flies have two PCGF proteins specific for canonical PRC1, encoded by genes located next to each other, and just one distinct PCGF protein incorporated in variant RING1 complexes (Kang *et al*., 2022). Leveraging the power of *Drosophila* genetics, we discovered that canonical PRC1 and variant RING1 complexes monoubiquitylate H2AK118 across distinct genomic regions. We found that the PCGF protein specific for variant RING1 complexes, which we named Siesta, controls larval locomotion independently of H2AK118 ubiquitylation and is dispensable for the epigenetic repression of *Drosophila* homeotic genes. Exploiting the division of labour between PRC1 and Siesta-RING1 complexes, we employed thousands of reporters integrated in parallel to conclude that H2AK118 ubiquitylation has no major repressive effect on transcription.

## RESULTS

To clarify the contribution of PRC1 and variant RING1 complexes to H2AK118 mono-ubiquitylation and the repression by the Polycomb system, we turned to the two loss-of-function alleles of the *l(3)73Ah* gene, which we briefly mentioned in (Kang *et al*., 2022). The PCGF protein encoded by the *l(3)73Ah* gene is an ortholog of the common ancestor of mammalian PCGF1, PCGF3 and PCGF5. It forms two distinct RING1-RYBP complexes that include either Kdm2, RSF1, BCOR, and SkpA or Tay and CKIIα/β (Kang *et al*., 2022). The loss-of-function alleles of the *l(3)73Ah* gene generated by CRISPR/Cas9-mediated genome editing contain 1052bp and 868bp deletions that remove the start codon and almost the entire open reading frame of the gene (Figure 1A, Figure S1). Animals trans-heterozygous for the two alleles die during pupation (Figure 1B), consistent with the reported phenotype of the original (now lost) ethyl methanesulfonate (EMS)-induced mutant allele (Belote et al., 1990; Irminger-Finger and Nothiger, 1995). Normally, *Drosophila* larvae continuously forage for food as they have a limited time to gain the weight required to undergo metamorphosis (Tyson et al., 2023). In contrast, the trans-heterozygous mutant larvae display an obvious locomotion defect (described in detail in a later section), which manifests in prolonged periods of inactivity between bouts of crawling. The lethality and the locomotion defect of the new loss-of-function mutants are complemented by one copy of the transgene containing 2.5kb fragment of genomic DNA that encompasses the entire *l(3)73Ah* transcript plus 613bp of the DNA upstream of the Transcription Start Site (TSS) and 129bp downstream of the transcript end (Figure 1A). This assures that the lethality and the locomotion defect are caused by the disruption of *l(3)73Ah* function and not due to a second-site mutation.

**Figure 1.**
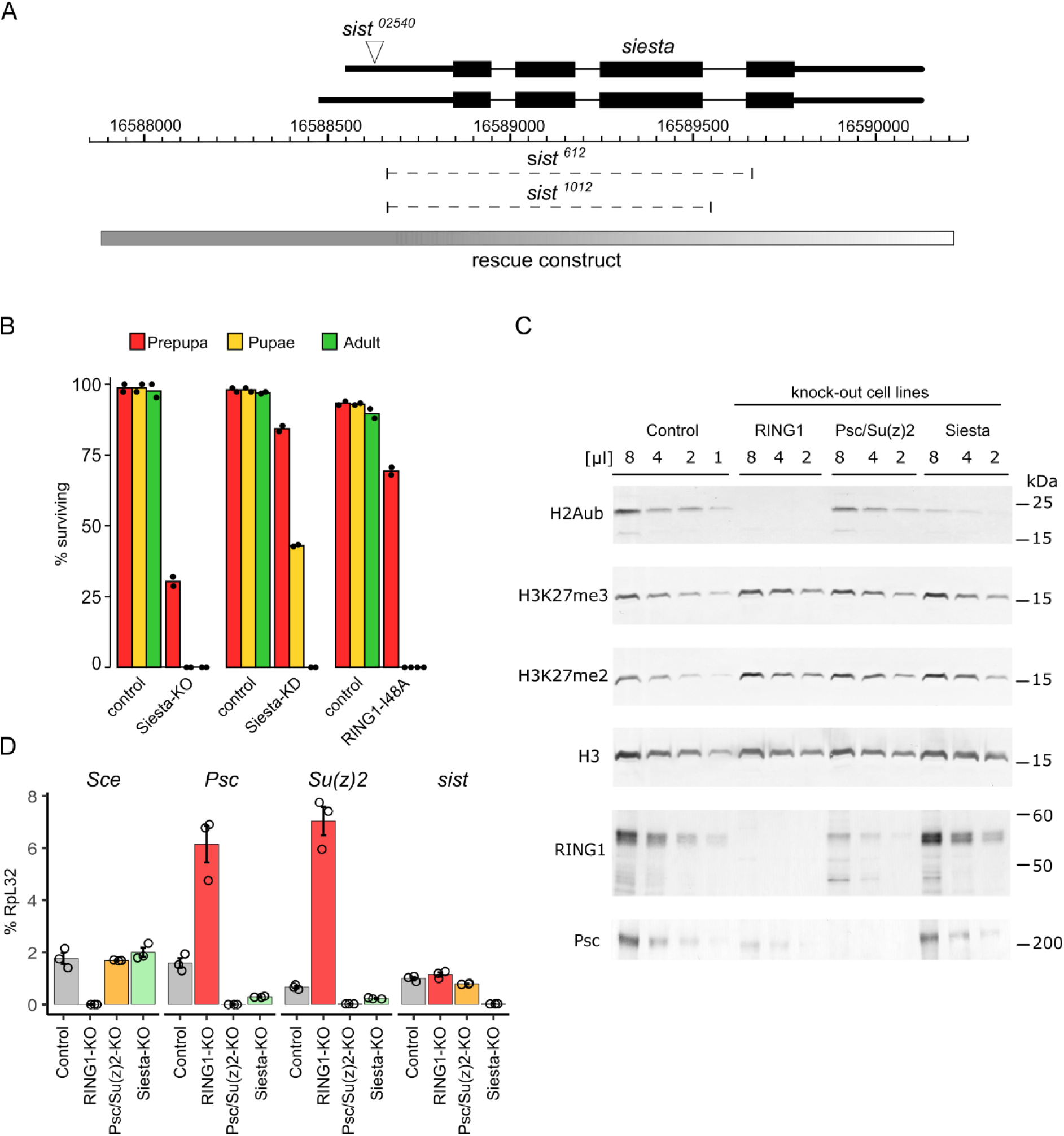
*siesta* alleles and the interrelation between PRC1 and Siesta-RING1 complexes. **A.** The schematics of the *siesta* locus. Two alternative transcripts above the coordinate scale (*dm6* genomic release) are shown with Transcription Start Sites (TSS) to the left. Thin lines indicate introns, and black boxes correspond to the coding parts. The position of transposon insertion in the *sist^02540^*allele (also known as *l(3)73Ah^02540^*) is indicated by a white triangle. Dashed lines mark the extent of *sist^612^* and *sist^1012^*deletions. The grey box indicates the genomic fragment sufficient to complement the *sist* loss-of-function mutations. **B.** The lethal stage of Siesta-KO (*sist^612^*/*sist^1012^*), Siesta-KD (*sist^612^*/*sis^02540^*) and RING1-I48A (*sce^I48A^*/*sce^I48A^*) mutants. Matching heterozygous controls were used for comparison. The bar plots indicate the average survival rate, starting with 150 first instar larvae. The black dots show the results of two independent experiments. **C.** Two-fold dilutions of total nuclear protein extracts from the control, RING1-KO, Psc/Su(z)2-KO and Siesta-KO cell lines were analysed by western blot with antibodies indicated to the left. Positions of molecular weight markers (in kilodaltons) are shown to the right. **D.** Relative transcript abundance for the *Sce*, *Psc*, *Su(z)2* and *sist* genes in the control, RING1-KO, Psc/Su(z)2-KO and Siesta-KO cells. The bar plots show the average of three independent RT-qPCR experiments performed with three independently prepared RNA samples. Circles show results of individual experiments, and whiskers indicate standard error of the mean.

The discoverers gave the original *l(3)73Ah* gene name to mark the gene’s position relative to other complementation groups identified in their mutagenic screen (Belote *et al*., 1990). This name has no connection to the gene function and is hard to remember, pronounce, and write. Following the long-standing tradition of naming *Drosophila* genes by their visible mutant phenotype, we propose to rename *l(3)73Ah* to *siesta* (*sist*) and will refer to the gene and its product as such hereafter. In line with the proposed nomenclature, we will refer to the loss-of-function alleles as *sist^612^* (formerly *l(3)73Ah^612^*) and *sist^1012^* (formerly *l(3)73Ah^1012^*).

### Multiple factors control the relative abundance of PRC1 and Siesta-RING1 complexes

We reasoned that a smaller repertoire of *Drosophila* PCGF proteins could provide a clearer picture of the genomic distribution and extent of H2AK118 ubiquitylation mediated by canonical PRC1 versus variant RING1 complexes. To avoid the problem of maternal contribution and to have unrestricted amounts of material, we derived a cultured cell line from *Drosophila* embryos homozygous for the *sist^612^*allele using the Ras^V12^ transformation approach (Simcox et al., 2008). Although *sist* is essential for viability, the mutant cells (hereafter referred to as Siesta-KO) are viable and proliferate in culture. Consistent with previous RNAi knock-down studies (Kahn *et al*., 2016), Siesta-KO cells lose approximately 80% of the bulk H2AK118ub, detected in the control Ras^V12^ transformed but otherwise wild-type cells (Figure 1C). This is a substantially greater reduction of the bulk H2AK118ub compared to ∼20% loss seen in Psc/Su(z)2-KO cells (Figure 1C). To ensure that the antibodies used to assay the H2AK118ub are specific, we edited the genome of Ras^V12^ transformed control cells (see Methods section for details) and derived a cultured cell line where the *Sce* gene encoding the RING1 protein is deleted. In these cells (hereafter referred to as RING1-KO), western blot analysis detects no trace of H2AK118ub, affirming that our assay is highly specific (Figure 1C). Although H2AK119ub was shown to stimulate the catalytic activity of human PRC2 *in vitro* (Kalb *et al*., 2014; Kasinath *et al*., 2021), we see no obvious reduction of the bulk di- or tri-methylated H3K27 in Siesta-KO or RING1-KO cells (Figure 1C).

The relation between intranuclear levels of RING1, Psc and Siesta appears more complicated. As expected, RING1 is undetectable in RING1-KO cells, and Psc is absent in Psc/Su(z)2-KO cells (Figure 1C). Surprisingly, we observed a substantially (∼4 times) lower amount of RING1 in Psc/Su(z)2-KO cells and, conversely, ∼4 times less Psc in RING1-KO cells (Figure 1C). On the other hand, the differences in the levels of RING1 and Psc in Siesta-KO cells, if any, were too small to be reliably detected (Figure 1C). As no antibodies against Siesta are currently available, we could not track its levels in our cultured cell lines. The RT-qPCR analysis indicated that transcription of the *Sce* gene is the same in the control, Psc/Su(z)2-KO and Siesta-KO cells (Figure 1D), indicating that the reduction in RING1 protein abundance occurs post-transcriptionally. Although we cannot formally exclude that Psc/Su(z)2-KO and Siesta-KO are needed for the efficient translation of RING1 mRNA, it is more plausible that RING1 becomes unstable when not in the complex with Psc, Su(z)2 or Siesta. Given that Psc/Su(z)2-KO has a more pronounced effect on nuclear RING1 pool than Siesta-KO, our data suggest that the majority of RING1 resides in the complex with Psc/Su(z)2.

The effects of RING1 and Siesta knock-out on the nuclear Psc pool are more complicated. *Psc* and *Su(z)2* genes are situated next to each other. The gene cluster includes multiple Polycomb Response Elements (PREs) bound by PRC1 and PRC2 (Park et al., 2012; Schwartz *et al*., 2006), resulting in a feedback loop where mutations in genes encoding for PRC1 subunits increase *Psc* and *Su(z)2* transcription (Ali and Bender, 2004). In agreement with this, *Psc* and *Su(z)2* transcription in RING1-KO cells increases four-to ten-fold (Figure 1D). Despite this, the amount of Psc protein in RING1-KO cells is notably lower than that in control cells, suggesting that Psc is unstable in the absence of RING1, and that elevated transcription of the *Psc* gene cannot compensate for this. Curiously, in Siesta-KO cells, transcription of the *Psc* and *Su(z)2* genes is reduced (Figure 1D). No anti-Su(z)2 antibodies are currently available, so we could not track it in our experiments. We speculate that, in *Drosophila* cells, the amount of RING1 is limited. In the absence of Siesta, more RING1 protein is available for incorporation into PRC1, which, in turn, increases the intranuclear concentration of PRC1 and leads to stronger transcriptional repression of the *Psc-Su(z)2* locus. Such a feedback loop may have evolved to keep the appropriate balance between PRC1 and Siesta complexes sharing the common RING1 subunit.

To conclude, our observations indicate that in *Drosophila* cells, most of the RING1 protein resides in complexes with Psc/Su(z)2. Yet, it is the comparatively less abundant Siesta-RING1 complexes that are responsible for generating the bulk of steady-state H2AK118ub.

### PRC1 and Siesta-RING1 complexes ubiquitylate H2AK118 in different parts of the genome

Sequencing of DNA immunoprecipitated with anti-H2AK118ub antibodies from chromatin lysates of control cells (H2AK118ub ChIP-seq) revealed a broad distribution of the ChIP-seq signal throughout the genome (Figure 2A). In contrast, H2AK118ub ChIP-seq with chromatin lysates from RING1-KO cells resulted in a uniform and low signal across the genome confirming that our assay is accurate and specific (Figure 2A). To enable quantitative comparison of ChIP-seq signals in cells with different genomic backgrounds, we adopted the “sans-spike-in” approach for quantitative ChIP-sequencing (siQ-ChIP) proposed by Dickson and colleagues (Dickson et al., 2023; Dickson et al., 2020). This approach is straightforward and avoids potential pitfalls associated with adding small amounts of chromatin from other species to the immunoprecipitation reaction (so-called “spike-in” approach) that is often used for ChIP-seq signal normalization (Dickson *et al*., 2020). Briefly, all ChIP-seq assays were performed with a fixed amount of crosslinked chromatin lysates and the same amount of a given antibody. For each sample, we recorded the fraction of immunoprecipitated DNA that was used for sequencing. The values were then used for scaling ChIP-seq signals to represent total ChIP yields (i.e. the signal expected if the entire immunoprecipitated DNA has been sequenced), adjusted for the sequencing depth (see the Methods section for a detailed description of the procedure). We performed two independent ChIP-seq experiments for each cell line and antibody and sequenced the DNA from corresponding chromatin input materials to control for potential sample processing and sequence alignment biases. The genomic distributions of ChIP-seq signals for the corresponding replicate experiments are highly concordant (r = 0.90 for control cells and 0.74 for RING1-KO cells, also see Table S1 and Figure S2A).

**Figure 2.**
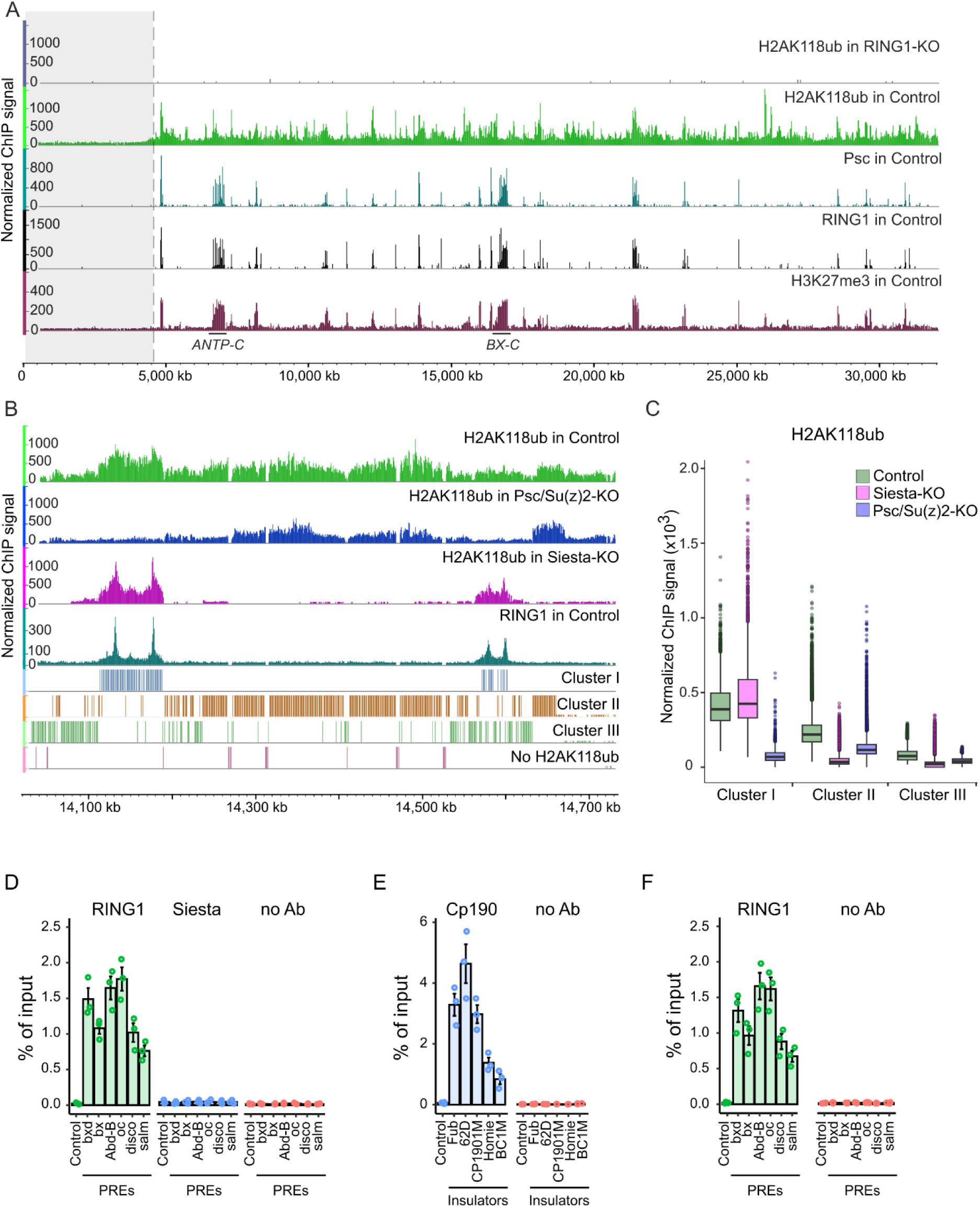
Genome-wide H2AK118 ubiquitylation by PRC1 and variant RING1 complexes. **A.** Genome browser tracks showing normalized ChIP-seq signals for H2AK118ub in RING1-KO and control cell lines across chromosome 3R. Tracks for Psc, RING1, and H3K27me3 in control cells are also included to pinpoint positions of genes repressed by the Polycomb system. The location of the Antennapedia complex (ANTP-C) and the bithorax complex (BX-C) is indicated by black lines above the coordinate scale (*dm6* genome release). The shaded region marks the extent of pericentromeric heterochromatin as defined in (Riddle *et al*., 2011). **B.** Normalized ChIP-seq signals for H2AK118ub and RING1 in control, Psc/Su(z)2-KO, and Siesta-KO cell lines across a representative region of chromosome 3L. The genome segmentation into 1kb regions of four different categories (Cluster I, Cluster II, Cluster III, and regions lacking H2AK118ub) is shown below the ChIP-seq signal tracks. **C.** H2AK118ub ChIP-seq signals in 1kb genomic regions of different categories in control, Siesta-KO, and Psc/Su(z)2-KO cells. Note dramatic reduction of the signal in Psc/Su(z)2-KO but not Siesta-KO cells at PRE-equipped loci (Cluster I) compared to control cells. At other regions (Cluster II and III), the loss of Siesta has a greater effect on H2AK118ub signal compared to Psc/Su(z)2-KO. Box plots indicate medians and span the interquartile range with whiskers extending 1.5 times the range. Dots designate the outliers. **D.** ChIP-qPCR with crosslinked chromatin lysates prepared from embryos where the loss of endogenous *siesta* function was complemented by transgenic expression of tagged Siesta protein (*sist::Twin-Strep-Myc-sist*; *sist^612^*/*sist^1012^* embryos) show robust immunoprecipitation of PREs with antibodies against RING1 but not with Strep-Tactin resin, specific for tagged Siesta protein. Here and in **E** and **F**, the bar plots show the average of three independent ChIP-qPCR experiments performed with three independently prepared chromatin samples. Circles show results of individual experiments, and whiskers indicate standard errors of the mean. Control experiments with chromatin from embryos that express Cp190 protein tagged with Twin-Strep-Tag show robust immunoprecipitation of insulator elements (Kahn et al., 2023) with Strep-Tactin resin (**E**) and PREs (Kahn *et al*., 2016) with anti-RING1 antibodies (**F**).

A side-by-side comparison of H2AK118ub ChIP-seq signals, averaged over 1kb genomic windows, for the control and RING1-KO cells indicates that 82% of the annotated *Drosophila* genome exhibit detectable levels of H2AK118ub. In most of the windows (78%) where the H2AK118ub was not detected, this happened because the windows contained repeated DNA and could not be uniquely matched to the sequencing reads. The remaining windows with unique nucleotide sequences but no detectable H2AK118ub are scattered throughout the genome, with notably higher density within pericentromeric regions. These regions, sometimes referred to as pericentromeric heterochromatin, are decorated with the HP1 protein and nucleosomes di- and tri-methylated at Lysine 9 of histone H3 (Riddle et al., 2011). Interestingly, even excluding the windows without detectable H2AK118ub, the windows corresponding to pericentromeric heterochromatin display substantially lower H2AK118ub ChIP-seq signals compared to the rest of the genome (Figure 2A, S2B-C). A similar observation was made during H2AK118ub profiling in spermatocytes and wing imaginal disc cells (Anderson et al., 2023). It is tempting to speculate that low H2AK118ub in the pericentromeric regions is common for many cell types due to HP1-mediated changes in chromatin structure that hinder the action of RING1 complexes.

To understand the contribution of PRC1 and Siesta-RING1 complexes to the H2AK118 ubiquitylation throughout the genome, we mapped the H2AK118ub distribution in Siesta-KO and Psc/Su(z)2-KO (Kahn *et al*., 2016) cells. We supplemented these experiments with ChIP-seq mapping of H3K27me3, Psc and RING1 in the control cells, which revealed sites where RING1 complexes are bound stably and marked the genes repressed by the Polycomb system, including positions of associated PREs. We also mapped RING1 in the RING1-KO cells to confirm the specificity of the anti-RING1 antibody.

Visual inspection of ChIP-seq signals shows that peaks of RING1 ChIP-seq signal (the sites of stable RING1 binding) are confined to peaks of Psc signal embedded in stretches of chromatin enriched in H3K27me3 (Figure 2A, S2A). This indicates that PREs are the only sites in the *Drosophila* genome stably bound by RING1 complexes. In line with this suggestion, stretches of chromatin around PREs with elevated ChIP-seq signals for H3K27me3 typically show elevated signals for H2AK118ub (Figure 2A, S2A). These H2AK118ub signals tend to be higher than those for chromatin fragments elsewhere in the genome (Figure S2D). However, there are notable exceptions. As pointed out earlier, the chromatin of the bithorax complex and Antennapedia homeotic gene clusters contains very little H2AK118ub (Bonnet *et al*., 2022; Lee *et al*., 2015), despite these genes being repressed and their PREs bound by RING1 (Figure S3). The systematic search for PRE-equipped loci with elevated H3K27me3 but low H2AK118ub revealed two similar cases: *β amyloid protein precursor-like* (*Appl)* – *ventral nervous system defective* (*vnd)* locus and *Visual system homeobox 1 (Vsx1)* - *Visual system homeobox 2 (Vsx2)* gene cluster (Figure S3, S4). Products of these four genes control aspects of nervous system development, and *vnd*, *Vsx1* and *Vsx2* encode homeobox transcription regulators (Erclik et al., 2008; Jimenez et al., 1995; Luo et al., 1990). As suggested by Bonnet and colleagues, the low steady-state levels of H2AK118ub at these loci may be explained by an exceptionally efficient de-ubiquitylation of H2AK118ub by the PRE-bound PR-DUB complex (Bonnet *et al*., 2022). We speculate that mutations in genes encoding PR-DUB subunits may have a particularly strong negative impact on the epigenetic repression of *Appl*, *vnd*, *Vsx1* and *Vsx2*, similar to that observed for homeotic genes of the bithorax complex (Scheuermann et al., 2010).

The H2AK118ub ChIP-seq signal at PRE-equipped loci is dramatically reduced in Psc/Su(z)2-KO cells but remains unaffected or even slightly increased in Siesta-KO cells (Figure 2B). This argues that H2AK118ub is catalysed by Psc/Su(z)2 -containing RING1 complexes, most likely canonical PRC1. Further strengthening this conclusion, we detected no Twin-Strep-tag-mediated immunoprecipitation of PREs DNA out of crosslinked chromatin lysates prepared from embryos where the loss of endogenous *siesta* function was complemented by transgenic expression of tagged Siesta protein (*sist::Twin-Strep-Myc-sist*; *sist^612^*/*sist^1012^* embryos) (Figure 2D-F). This indicates that, in contrast to PRC1, Siesta-RING1 complexes are not attracted to PREs.

As estimated previously (Lundkvist et al., 2023), regions surrounding PREs that are highly enriched in H3K27me3 and decorated by PRC1-dependent H2AK118ub account for only ∼ 5% of the *Drosophila* genome. Most of the remaining genome also shows significantly elevated H2AK118ub ChIP-seq signal, with some of the sites displaying signals comparable in magnitude to those seen at PRE-equipped loci (Figure 2A, S2A). None of these sites display elevated ChIP-seq signals for RING1, suggesting that H2AK118 ubiquitylation outside PRE-equipped loci is installed without stable binding of RING1 complexes. Importantly, this widespread H2AK118 ubiquitylation is strongly reduced in Siesta-KO cells (Figure 2B, S2A). The effect is not limited to cultured cells. The immunostaining of polytene chromosomes from the *sist^02540^*/*sist^612^*(Sist-KO/KD) third instar larvae shows equally dramatic loss of H2AK118ub throughout chromosome arms except at PRE-equipped loci, most brightly stained with antibodies against H3K27me3 (Figures S2E-F).

Unsupervised k-means clustering of 1kb genomic segments (bins) based on comparison of H2AK118ub ChIP-seq signals in the control, Psc/Su(z)2-KO, and Siesta-KO cells reinforces the observations above. The clustering partitions the genome into regions of three kinds: PRE-equipped loci with high H2AK118ub dependent on canonical PRC1 (Cluster I), regions with high H2A118ub strongly dependent on Siesta-RING1 complexes (Cluster II), and regions with low but significant H2A118ub also primarily dependent on Siesta-RING1 complexes (Cluster III). Completing the segmentation of the genome are regions devoid of H2AK118ub discussed above and excluded from the unsupervised k-means clustering procedure (Figure 2B, Figure S5). Curiously, a side-by-side comparison of H2AK118ub ChIP-signals in Cluster II and Cluster III regions in Psc/Su(z)2-KO and control cells reveals a small but detectable reduction of the signal upon Psc/Su(z)2 loss (Figure 2C). This argues for modest hit-and-run H2AK118 ubiquitylation activity of untethered PRC1 or Psc/Su(z)2-containing variant RING1 complexes genome-wide.

To summarize, our observations indicate that PREs are the only sites in the *Drosophila* genome stably bound by RING1 complexes. These complexes, most likely canonical PRC1, contain Psc or Su(z)2 proteins and produce most of the H2AK118ub at PRE-regulated genes. At the same time, the hit-and-run action of Siesta-RING1 complexes is responsible for the bulk of H2A118ub elsewhere in the genome.

### Siesta complexes control larval locomotion independently of H2AK118 ubiquitylation

Trans-heterozygous *sist^612^*/*sist^1012^*zygotic null larvae do not exhibit homeotic transformations but display a pronounced locomotion defect characterized by reduced crawling speed. This phenotype is distinct from that of the mutants for genes encoding canonical PRC1 subunits. Given that Siesta-RING1 complexes monoubiquitylate H2AK118 outside the genes controlled by PRC1, we asked whether the locomotion defects observed in *sist* mutants are linked to the widespread depletion of H2AK118ub across the genome.

To this end, we tracked the locomotion of *sist^612^*/*sist^1012^*(Siesta-KO) second instar larvae and compared them to the locomotion of the *sist^612^*/*sist^02540^* (Siesta-KD) and *Sce^I48A.gen^; Sce^KO^* (RING1-I48A) animals. The *sist^02540^* allele is the insertion of the *PZ* transposable element (Spradling et al., 1999) in the 5’UTR of *sist* (Figure 1A). As shown in Figure 1B, *sist^612^*/*sist^02540^*animals survive longer than Siesta-KO, indicating that *sist^02540^* retains partial function and should be considered as a hypomorphic allele. The RING1-I48A animals are homozygous for the loss-of-function *Sce^KO^* allele (Gutierrez et al., 2012) and carry a transgene expressing a catalytically impaired RING1 protein in which isoleucine 48 is substituted with alanine (Figure S6). This mutation disrupts RING1’s E3 ligase activity toward H2AK118 (Pengelly *et al*., 2015). Like Siesta-KO mutants, RING1-I48A animals die during pupation (Figure 1B).

The Siesta-KO mutants move 5 times slower than heterozygous control larvae (Figure 3A). The hypomorph Siesta-KD larvae move faster than Siesta-KO, and the RING1-I48A display the highest median speed of the three (Figure 3A). Strikingly, these locomotion phenotypes did not correlate with the extent of bulk H2AK118ub loss (Figure 3C). The latter is the most pronounced in the RING1-I48A larvae and the least affected in Siesta-KD mutants. These findings suggest that Siesta complexes control larval locomotion through a mechanism that is independent of H2AK118 ubiquitylation.

**Figure 3.**
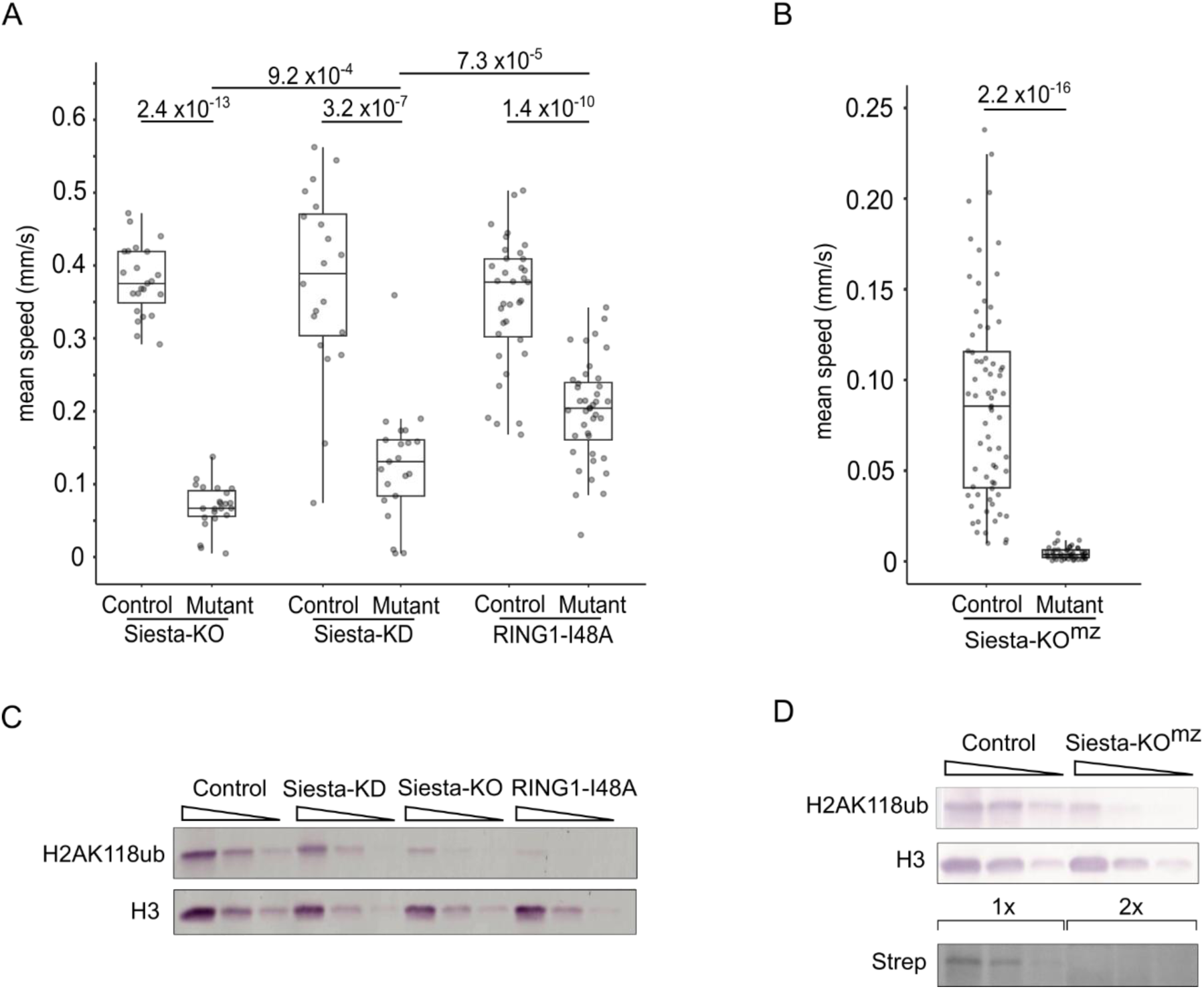
Siesta complexes control locomotion independently of H2AK118 ubiquitylation. **A.** Motion tracking of Siesta-KO (*sist*^612^/*sist*^1012^), Siesta-KD (*sist*^612^/*sist*^02540^) and RING1-I48A (*Sce*^148A.gen^; *Sce*^KO^) second instar larvae. Heterozygous animals were used as matching controls. Here and in **B,** the box plots show median speeds and span the interquartile range with whiskers extending to cover all observations. Grey dots indicate the mean speeds of individual larvae. The differences in medians between corresponding groups were tested for statistical significance using the Wilcoxon rank sum test, and p-values are displayed above the box plots. **B.** Motion tracking of Siesta-KO^mz^ and matching control first instar larvae. **C.** Western blot analysis of H2AK118ub levels in various *siesta* mutants. Three-fold serial dilutions of total protein extract from second instar larvae were assayed using anti-H2AK118ub and pan-histone H3 (loading control) antibodies. **D.** Three-fold serial dilutions of total protein extract from Siesta-KO^mz^ and control embryos, where the transgenic copy of Siesta was not removed, were analysed by western blot with anti-H2AK118ub and anti-H3 antibodies. The Twin-Strep-tagged transgenic Siesta was detected using the Streptavidin-AP conjugate. Note that, in Siesta-KO^mz^ embryos, the transgenic Siesta protein is not detected even when twice as much protein extract is used compared to the control.

Many *Drosophila* regulators, including the Polycomb group proteins, are maternally supplied. In such cases, zygotic loss-of-function mutants derived from heterozygous parents begin development with maternally deposited wild-type mRNA and protein, which is often sufficient to support the early development and partially mask mutant phenotypes. To understand the role of maternally contributed Siesta, we used the strain where *sist^612^*/*sist^1012^* mutations were initially rescued by a transgenic *sist* copy, which was subsequently excised specifically in the germline using FLP recombinase (Figures S7, S8). This approach generated Siesta-KO^mz^ mutants that lack both maternally deposited and zygotically expressed Siesta. The absence of Siesta protein and substantially reduced H2AK118ub in 0-24 hour-old embryos are evident from the western blot analysis (Figure 3D). None of the Siesta-KO^mz^ mutants progressed to the second instar larval stage, and the first instar larvae exhibited severely impaired locomotion compared to their zygotic counterparts (Figure 3B, Supplementary movies 1, 2).

To assess whether the reduced locomotion was due to general sensory impairment, we tested the larvae’s response to hypoxia. In *Caenorhabditis elegans*, reduced oxygen levels trigger increased forward movement as an escape response (Onukwufor et al., 2022; Zhao et al., 2022). Similarly, both control and Siesta-KO^mz^ larvae increased their crawling speed when exposed to 1% oxygen, although the mutants remained significantly slower (Figure S9, Supplementary movies 3, 4). These results suggest that the locomotor defect in *siesta* mutants is not caused by general sensory deprivation. The precise nature of the defect warrants further neuro-anatomical investigation.

### *Drosophila* RYBP is not essential for viability and bulk H2AK118 ubiquitylation

In mammals, the presence of the RYBP subunit, or its paralogue YAF2, distinguishes variant RING1/RING2 complexes from canonical PRC1, which instead incorporates one of the CBX subunits (Gao *et al*., 2012; Wang et al., 2010). The same applies to *Drosophila*, which encodes a single RYBP orthologue (Kang *et al*., 2022). The simplicity of the *Drosophila* system, with only one RYBP gene, provides an opportunity to dissect its functional contribution to RING1 complexes. Flies homozygous for *RYBP^KG08683^*, the only allele designated as loss-of-function in the literature, display reduced viability, substantial developmental delay and 90% female sterility (Gonzalez et al., 2008). However, the *RYBP^KG08683^* allele is caused by a P-element insertion in the 5’-UTR of *RYBP*, raising the possibility that it does not fully abolish gene function.

To address this uncertainty, we generated new *RYBP* mutations using CRISPR/Cas9-mediated genome editing. To this end, we designed guide RNAs targeting the Cas9 endonuclease to the 5’ UTR and 3’UTR of the *RYBP* transcript (Figure 4A). As a result, we recovered two new alleles of the *RYBP* gene, which remove 880bp (*RYBP^4-2^*) and 902bp (*RYBP^5-1^*) and thus eliminate the entire open reading frame (ORF) (Figure 4A, Figure S10). To our surprise, and in contrast to Siesta-KO, these clean *RYBP* loss-of-function mutants (RYBP-KO) are homozygous viable, fertile, and can be maintained as stable stocks. This suggests that the developmental defects observed in *RYBP^KG08683^*flies are due to a second-site mutation or that the allele functions as a gain-of-function rather than a true null. Western blot analysis revealed no difference in bulk H2AK118ub levels between RYBP-KO and control flies (Figure 4B), indicating that RYBP is not essential for global H2AK118 ubiquitylation in *Drosophila*.

**Figure 4.**
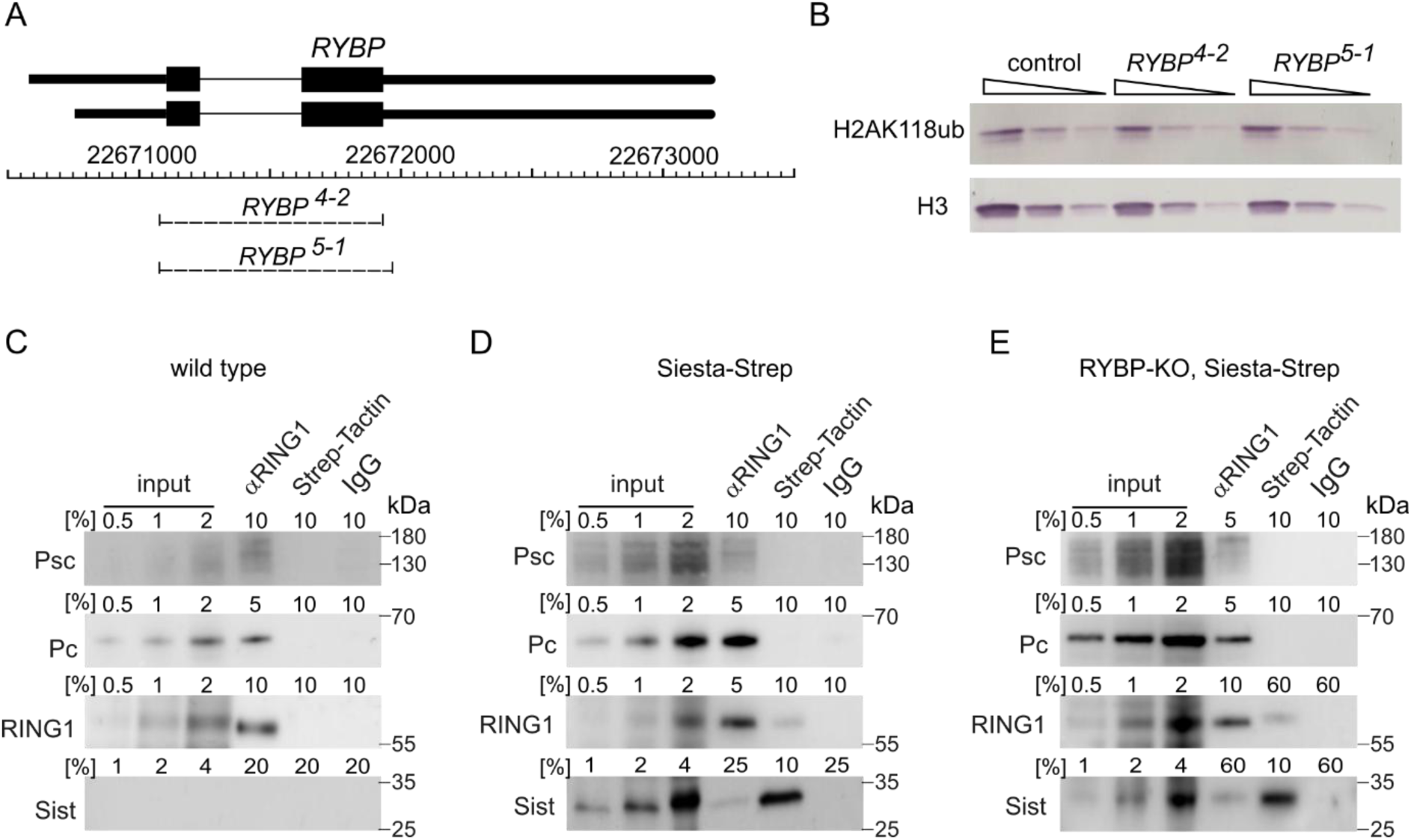
Drosophila RYBP is dispensable for the bulk of H2AK118 ubiquitylation. **A.** The schematics of the *RYBP* locus. Two alternative transcripts above the coordinate scale (*dm6* genomic release) are shown with Transcription Start Sites (TSS) to the left. Thin lines indicate introns, and black boxes correspond to the coding parts. Dashed lines mark the extent of *RYBP^4-2^*and *RYBP^5-1^* deletions. **B.** Three-fold dilutions of total protein extract from wild-type control, *RYBP^4-2^* and *RYBP^5-1^* second instar larvae were analysed by western blot with antibodies against H2AK118ub and H3 (loading control). The assay shows no reduction of H2AK118ub in animals lacking RYBP. The co-immunoprecipitation of Psc, Pc, RING1 and Sist from total nuclear protein extracts of wild type (**C**), Siesta-Strep (**D**) and RYBP-KO, Siesta-Strep embryos (**E**). The indicated fractions of material immunoprecipitated with anti-RING1, Strep-Tactin and total rabbit IgG (negative control) were analysed by western blot with antibodies indicated to the left of each panel. Strep-Tactin-HRP conjugate was used to detect the Twin-Strep-tagged transgenic Siesta protein. Positions of molecular weight markers (in kilodaltons) are shown to the right of each panel. Note that Siesta and RING1 co-immunoprecipitate even when RYBP is absent (**E**), although the signals are much weaker and require a larger fraction of precipitated material to be detected.

To investigate the impact of RYBP loss on Siesta complexes, we generated the fly strain homozygous for RYBP-KO and Siesta-KO alleles complemented with a transgenic copy of *sist* expressing a fully functional Twin-Strep- and Myc-tagged Siesta protein (*RYBP^4-2^*, *sist:Strep-Myc-sist*; *sist^1012^* referred to as RYBP-KO, Siesta-Strep). We then compared the co-immunoprecipitation of RING1 with Siesta, Psc and Pc from nuclear extracts of RYBP-KO and Siesta-KO embryos and that from nuclear extracts of Siesta-KO embryos complemented with the same *sist* transgene as above (*sist:Strep-Myc-sist*; *sist^1012^*, referred to as Siesta-Strep). As illustrated by figures 4C-E, the co-immunoprecipitation of RING1 with Pc or Psc is not altered by the RYBP loss. In contrast, in the absence of RYBP, the co-immunoprecipitation of RING1 and Siesta is drastically (> 6-fold) reduced, although it is still detectable (Figures 4C-E).

The Siesta-KO cells lose approximately 70% of the bulk H2AK118ub (Kahn *et al*., 2016), Figure 1C) and it is well established that RING1 requires a PCGF partner for efficient ubiquitylation of H2AK118ub (Ben-Saadon et al., 2006; Bentley et al., 2011; Buchwald et al., 2006; Li et al., 2006; McGinty et al., 2014). Yet we see no reduction in the bulk H2AK118ub levels in the RYBP-KO flies despite the substantial loss of co-immunoprecipitation between RING1 and Siesta when RYBP is absent. The latter suggests that the loss of RYBP does not affect the assembly of the Siesta-RING1 complex *in vivo*. We posit that, without RYBP, the complex becomes less stable and easily dissociates during protein extract preparation, resulting in substantially lower co-immunoprecipitation. The increased stability conferred by RYBP may explain the higher E3 ligase activity of RYBP-RING2 complexes towards H2AK119 *in vitro* (Gao *et al*., 2012; Rose *et al*., 2016).

### Epigenetic repression of developmental genes works without Siesta

The Siesta-KO^mz^ larvae display no obvious homeotic transformations, suggesting that, in contrast to PRC1 subunits, Siesta is not essential for epigenetic repression of developmental genes. To further test this conjecture, we compared the expression of the developmental regulators *prospero* (*pros*), *Abdominal-B* (*Abd-B*) and *Antennapedia* (*Antp*) in Siesta-KO^mz^, Psc/Su(z)2-KO (animals homozygous for the *Su(z)2-1.b8* deletion produced by heterozygous parents) and control embryos. *Antp* and *Abd-B* are homeotic selector genes that encode transcription regulators necessary to specify the identity of embryonic parasegments (PS) 4-5 and 10-14, respectively. The *pros* gene is expressed in a set of cells in the central nervous system, specifying their fate. The correct spatial expression pattern of *Antp*,

*Abd-B* and *pros* is achieved by the competing action of transcriptional enhancers and epigenetic repression by the Polycomb system. As expected, embryos lacking a zygotic supply of the Psc and Su(z)2 proteins show wide misexpression of *Abd-B*, *Antp* and *Pros* in the nervous system (Figure 5). In contrast, the spatial expression patterns of *Abd-B*, *Antp* and *Pros* in the Siesta-KO^mz^ do not differ from those in the control embryos (Figure 5). These results indicate that Siesta complexes are not generally required for the epigenetic repression of *Drosophila* developmental genes.

**Figure 5.**
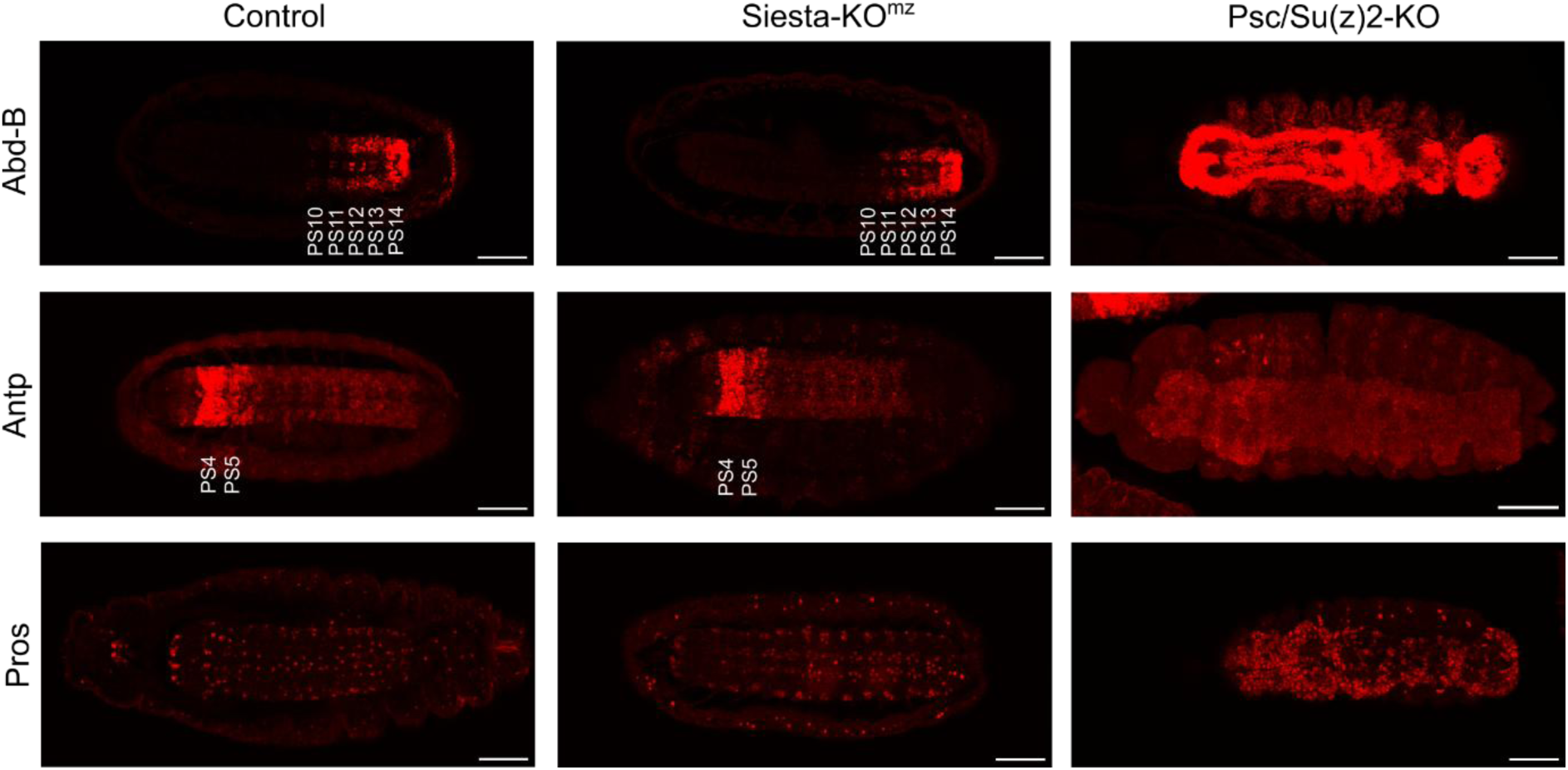
Siesta mutants maintain correct expression patterns of developmental genes. *Abd-B*, *Antp* and *Pros* expression was assayed by immunostaining of late-stage *Drosophila* embryos with corresponding antibodies. The embryo genotypes (Control, Siesta-KO^mz^ and Psc/Su(z)2-KO) are indicated above the image panel. A single optical section is shown for representative embryos stained with anti-Abd-B antibodies. Maximum intensity projection (MIP) of five Z stacks is shown for embryos stained with anti-Antp antibodies, and MIP of two Z stacks is shown for embryos stained with anti-Pros antibodies. Parasegments 10 to 14 (PS10-PS14) and 4 and 5 (PS4 and PS5) are indicated to highlight the *Abd-B* and *Antp* expression patterns in the control and Siesta-KO^mz^ embryos. Scale bars at the bottom right of each image correspond to 50µm.

### No evidence of H2AK118ub repressing transcription

Studies in mouse embryonic stem cells suggested that H2AK119 ubiquitylation (H2AK118ub in *Drosophila*) acts as a generic transcriptional repressor and a primary means by which the Polycomb system represses target genes (Blackledge and Klose, 2021). Contradicting this view, Müller and colleagues did not detect phenotypes characteristic of Polycomb group mutants in *Drosophila* deficient for H2AK118 ubiquitylation (Pengelly *et al*., 2015). Furthermore, transcriptome analyses of the mutants failed to detect any genes whose transcription significantly changed. The conventional RNA sequencing assay used in the study was designed to detect differences in the transcriptional output of a small subset of genes. Given the widespread distribution of H2AK118 ubiquitylation, its loss could have affected many genes at once, complicating the detection.

To overcome this limitation, we employed the Thousand Reporters Integrated in Parallel (TRIP) assay (Akhtar et al., 2013), which enables genome-wide assessment of transcriptional activity by measuring how the same reporter gene behaves when integrated at different genomic locations. If H2AK118ub functions as a strong repressor, reporter constructs inserted into regions with high H2AK118ub—such as PRE-associated loci (Cluster I) or regions enriched in Siesta-dependent H2AK118ub (Cluster II)— should exhibit lower transcriptional output compared to insertions in regions with low H2AK118ub (Cluster III). The division of labour between PRC1 and Siesta-RING1 complexes provides for an additional test. If the model is correct, the transcription of reporter genes integrated into Cluster I should be substantially higher in Psc/Su(z)2-KO cells compared to that in the control cells.

Conversely, reporters integrated in Cluster II regions should be transcribed more in Siesta-KO cells. Furthermore, insertions in both kinds of regions should produce more RNA in RING1-KO cells where H2AK118ub is globally depleted.

To test these predictions, we generated four TRIP libraries consisting of tens of thousands of reporter constructs flanked by inverted repeats from the *piggyBac* mobile element (Figure 6A). All libraries contained an identical *GFP* reporter gene fused at the 3’ end to an 18bp DNA segment with a random nucleotide sequence (barcode) (Figure 6A, B). In two of the libraries, the *GFP*-reporter was placed under the control *Hsp70Bb* gene promoter (Pelham, 1982; Xiao and Lis, 1988), while in the two other libraries, the *GFP* transcription was controlled by the *Metallothionein A* (*MtnA*) gene promoter (Bunch et al., 1988). Each library was distinguished by a specific 5bp sequence (promoter index). The *Hsp70Bb* promoter can drive robust medium-level transcription of chromatin-integrated reporters due to its open chromatin architecture that favours RNA Pol II recruitment (Shopland et al., 1995; Tsukiyama et al., 1994). The behaviour of the *MtnA* promoter in reporters integrated into the genome is less studied, but the transcription from the *MtnA* promoter could be induced by supplementing the cell culture growth medium with copper ions (Bunch *et al*., 1988).

**Figure 6.**
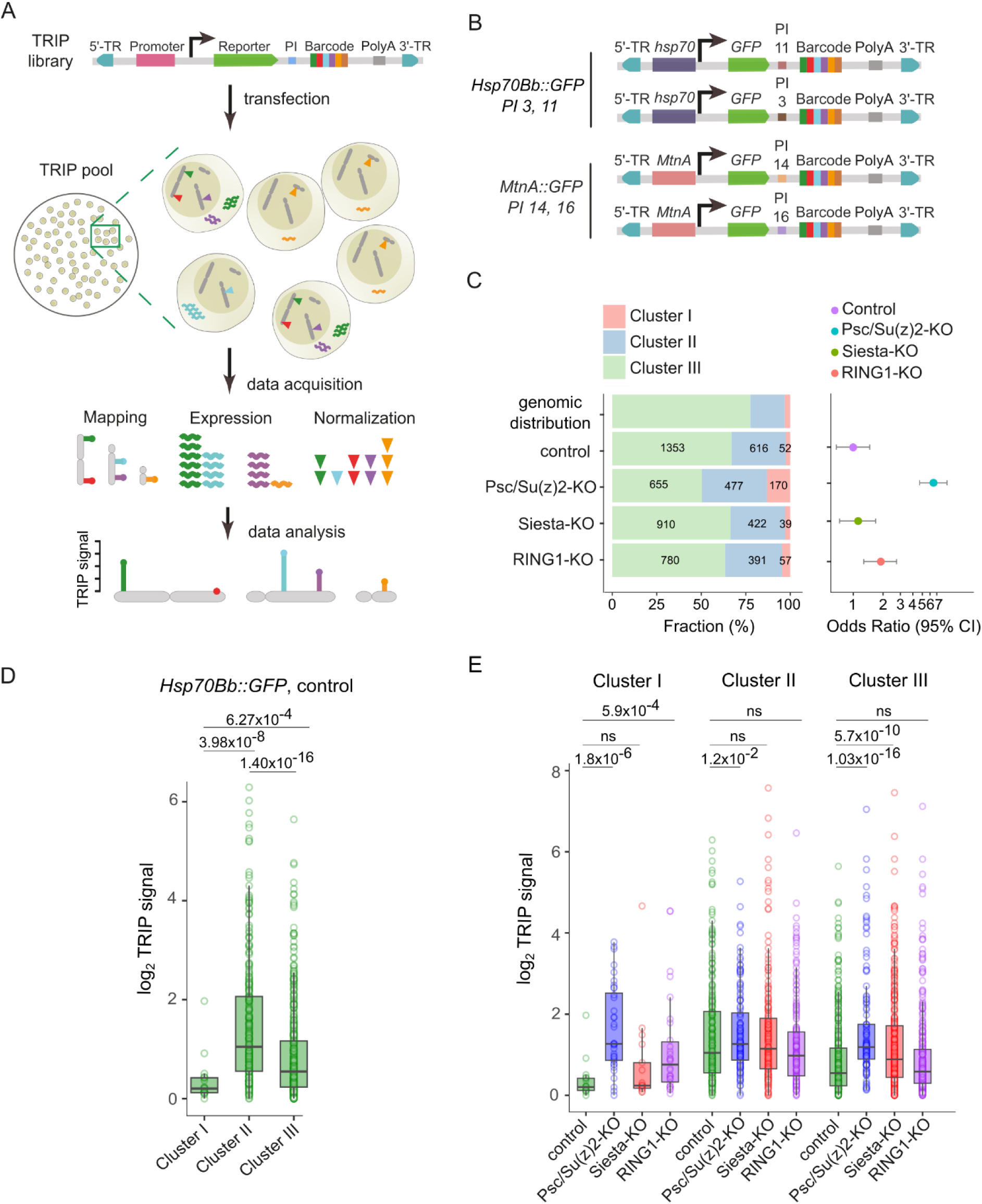
TRIP reveals no repressive effect of H2AK118 ubiquitylation on transcription. **A.** The schematic of the TRIP assay. The same colour code is used to illustrate reporter constructs integrated at distinct genomic sites and the corresponding RNA products. **B.** The schematic representation of the four TRIP libraries used for the assay. All constructs were integrated in the genome via piggyBac 5′ (5’-TR) and 3′ (3’-TR) terminal repeats, but each library was marked with a distinct promoter index (PI). **C.** Stacked bar plots show fractions and numbers of transgenic insertions within Cluster I-III regions in different cell lines compared to the relative abundance of each region type within the *Drosophila* genome. The dot plot to the left indicates the odds ratios of integration within PRE-regulated genes compared to the rest of the genome within different cell lines. Whiskers mark the 95% confidence interval. Note a significant increase in the odds of integrating into the PRE-regulated genes in the Psc/Su(z)2-KO cells. **D.** TRIP signals for the *Hsp70Bb::GFP* transgenes integrated in three types of genomic regions in control cells. Here and in **E**, the boxplots indicate the median and span interquartile range with whiskers extending 1.5 times the range and outliers shown as circles. The differences in medians between corresponding groups were tested for statistical significance using the Wilcoxon rank sum test, and p-values are displayed above the box plots. **E.** TRIP signals for the *Hsp70Bb::GFP* transgenes in different cell lines.

An approximately equimolar mixture of all four TRIP libraries was transfected into control, Psc/Su(z)2-KO, Siesta-KO and RING1-KO cells along with the plasmid expressing piggyBac transposase. The resulting pools of transgenic cells (TRIP cell pools) were used to map the genomic location of barcoded insertions by high-throughput sequencing of inverse PCR products. The transcriptional activity of corresponding transgenes in each TRIP cell pool, grown on media with and without copper, was deduced by counting barcodes after sequencing RNA produced by transgenes. To account for variations in the growth rates of individual cells in a TRIP cell pool and potential biases during sequencing library preparations, the barcode counts were further normalized by counting the barcode sequences within genomic DNA isolated from corresponding TRIP cell pools (Figure 6A, for further details, see the Methods section). All assays were performed in duplicates for each TRIP cell pool, starting from independent cell cultures.

Across all promoter variants and indices, we analysed 1228 to 2021 distinct reporter insertions per genetic background (Figure 6C). In all cases, the largest fraction of the insertions occurred within regions with low H2A118ub (Cluster III regions), followed by regions with high H2A118ub produced by Siesta-RING1 complexes (Cluster II regions) and a smaller number of insertions into the PRE-equipped loci (Cluster I regions). This distribution largely reflects the relative genomic representation of Clusters I–III (Figure 6C). Interestingly, we noted 7-fold higher odds of insertions into PRE-equipped loci (Cluster I regions) in Psc/Su(z)2-KO cells compared to control, Siesta-KO and even RING1-KO cells. This suggests that the PRE-equipped loci hinder the integration of *piggyBac*-based transgenes. This hindrance is dependent on Psc and/or Su(z)2 but not on H2AK118 ubiquitylation.

Analysis of TRIP signals from *Hsp70Bb::GFP* transgenes integrated into the genome of control cells shows that the transgenes integrated into PRE-equipped loci (Cluster I regions) are transcribed less than those integrated into Cluster II and Cluster III regions (Figure 6D). This is expected and agrees with previous observations that P-element-based transgenes become repressed by the Polycomb system when integrated close to PREs (Galloni et al., 1993; Park *et al*., 2012). These results confirm that our TRIP assay is sufficiently sensitive to detect the repressive effect of the Polycomb system.

Surprisingly, and contrary to the model in which H2AK118ub acts as a strong transcriptional repressor, the transgenes integrated into regions with high H2AK118ub (Cluster II, Figure 6D, Figure 2C) are more transcriptionally active compared to transgenes integrated into regions with lower H2AK118ub (Cluster III, Figure 6D, Figure 2C).

Consistent with the observations above, the transgenes integrated into PRE-equipped loci (Cluster I regions) show significantly higher transcriptional activity in Psc/Su(z)2-KO and RING1-KO cells compared to that in the control and Siesta-KO cells (Figure 6E). This reinforces the argument that transgenes integrated in these regions are repressed by the Polycomb system and that our TRIP assay is sufficiently sensitive to detect the repression. Contrary to the model proposing H2AK118ub as a transcriptional repressor, transgenes integrated in Cluster II and III regions are transcribed at the same level in RING1-KO cells, completely devoid of this modification (Figure 6E). In line with our previous findings that ablation of Psc and Su(z)2 leads to a weak but significant increase in transcription everywhere in the genome (Lee *et al*., 2015), *Hsp70Bb::GFP* transgenes integrated into Cluster II and III regions display higher transcription in Psc/Su(z)2-KO cells (Figure 6E). This effect is more pronounced for the generally lower-expressing insertions in Cluster III regions and does not correlate with the extent of H2AK118ub loss. The latter is significantly greater in Siesta-KO cells, in which transcription is essentially the same for insertions in Cluster II regions and just slightly elevated for insertions in Cluster III regions compared to control cells (Figure 6E).

The examination of TRIP signals from *MtnA::GFP* transgenes leads to similar conclusions. These transgenes display generally low transcription when cells are grown in the media without Cu^2+^ ions (Figure S11A). Nevertheless, the transcription of transgenes integrated into PRE-equipped loci (Cluster I regions) is weaker compared to those integrated elsewhere (Figure S11A). The TRIP signals from our *MtnA::GFP* transgenes increase when cells are grown on media supplemented with copper (Figure S11B, C). The median increase is lower than anticipated from the original study with transiently transfected constructs (Bunch *et al*., 1988), suggesting that the *MtnA* promoter is far less efficient in inducing transcription of chromatin-integrated transgenes. Even in the presence of Cu^2+^ ions, the transgenes integrated into PRE-equipped loci (Cluster I regions) are transcribed significantly less compared to those integrated elsewhere. Yet, we see no difference between *MtnA::GFP* transgenes integrated into Cluster II and Cluster III regions despite the former displaying significantly higher H2AK118ub ChIP-seq signals (Figure S11C). Like *Hsp70Bb::GFP* transgenes, the *MtnA::GFP* transgenes integrated into PRE-equipped loci (Cluster I regions) show significantly higher transcriptional activity in Psc/Su(z)2-KO and RING1-KO cells compared to that in the control and Siesta-KO cells (Figure S11D). On the other hand, the effects of Psc/Su(z)2, Siesta and RING1 knock-out on the transcription of transgenes integrated away from PREs are weak and variable. Importantly, we see no correlation between the extent of transcription and the loss of H2AK118ub.

To summarise, our TRIP assay demonstrates that, regardless of the promoter, transgenes integrated in the PRE vicinity tend to transcribe less than transgenes integrated elsewhere. The apparent transcriptional repression, which we attribute to the action of the Polycomb system, is relieved by removal of Psc/Su(z)2 or RING1 but not Siesta. However, the transgenes not influenced by the Polycomb system (i.e. integrated away from PREs) display no correlation between transcriptional activity and the degree of H2AK118ub within its integration site. Likewise, they show no consistent transcriptional changes correlated to the loss of H2AK118ub in various mutant backgrounds.

## DISCUSSION

Epigenetic repression of developmental genes requires canonical PRC1, which acts by interacting with H3K27me3 to propagate the repressed state of regulated genes. Canonical PRC1 is also an E3 ligase that can deposit a single ubiquitin molecule on K118/119 of histone H2A. This enzymatic activity is shared with other complexes formed by the RING1/RING2 subunit of canonical PRC1, although the latter cannot bind H3K27me3. It has been proposed that H2A ubiquitylation is an essential part of the repressive mechanism and that variant RING1/RING2 complexes evolved from ancestral canonical PRC1 to diversify the Polycomb system and enable the evolution of vertebrate-specific traits (Blackledge and Klose, 2021). Yet, systematic tracing of genes encoding core components of RING1/RING2 complexes in genomes from diverse animal clades has shown that the RING1/RING2 complexes found in vertebrates appeared early in animal evolution and likely served functions unrelated to epigenetic repression (Gahan *et al*., 2020). Our findings strongly support this view. In *Drosophila*, we show that canonical PRC1 and variant RING1 complexes monoubiquitylate H2A across distinct genomic regions. Furthermore, Siesta, the sole *Drosophila* PCGF protein specific for variant RING1 complexes, is dispensable for epigenetic repression of homeotic genes and controls larval locomotion independently of H2A ubiquitylation. Exploiting the division of labour between PRC1 and Siesta-RING1 complexes, we used the TRIP assay to demonstrate that H2A ubiquitylation has no major repressive effect on transcription. Below, we discuss the implications of our findings in more detail.

In the current literature, RING1/RING2 complexes are customarily referred to as variants of PRC1, providing for convenient nomenclature (e.g. PRC1.1, PRC1.2, etc). However, this terminology implicitly suggests that all RING1/RING2 complexes contribute to epigenetic repression. Misled by this nomenclature, a recent study considered RING1/RING2 complexes as a functionally coherent group and reported no coupling in the evolution of the PRC1 and PRC2 parts of the Polycomb system (de Potter *et al*., 2023). We believe this inference is premature. Instead, the co-evolution of canonical PRC1 and PRC2 should be examined independently of other RING1/RING2 complexes, as these two complexes are more clearly linked by their shared role in H3K27me3-mediated gene repression. To prevent similar misconceptions in the future, we suggest revising the current nomenclature and reserving the Polycomb Repressive Complex 1 name for canonical PRC1.

Consistent with earlier observations (Fursova *et al*., 2019; Lee *et al*., 2015), we find that most of the non-repeated *Drosophila* genome is packaged into nucleosomes monoubiquitylated at H2AK118.

Strikingly, the bulk of this widespread H2AK118 ubiquitylation is achieved without stable binding of RING1 complexes. In fact, PREs seem to be the only sites in the *Drosophila* genome stably bound by RING1 complexes. These complexes, likely canonical PRC1, contain Psc or Su(z)2 proteins and produce most of the H2AK118ub at PRE-regulated genes. How PREs tether PRC1 is not well understood. According to one model, PREs are comprised of binding sites for multiple DNA-binding proteins that combine individually weak interactions with PRC1 to yield the robust tethering (Schwartz, 2017). Which of the PRC1 subunit(s) interact with these DNA-binding proteins is not known, but our observations suggest it is not RING1. Otherwise, Siesta-RING1 complexes would also be bound to PREs.

Changes of RING1 levels in Siesta-KO and Psc/Su(z)2-KO cells indicate that most of the *Drosophila* RING1 protein resides in the complex with Psc/Su(z)2. Yet, the hit-and-run action of less abundant Siesta-RING1 complexes produces most of the steady-state H2AK118ub. This suggests that Siesta-RING1 complexes are more active E3 ligases. Stimulation by the RYBP subunit (Gao *et al*., 2012; Rose *et al*., 2016) or the differences in the biochemical properties of distinct PCGF proteins (Teslenko and Fierz, 2025) were proposed as explanations of the increased activity of variant RING1/2 complexes. Our genetic data support the latter view. Flies lacking the RYBP function are viable, fertile and show no detectable differences in the bulk H2AK118ub, although the RYBP loss results in substantially lower recovery of Siesta-RING1 complexes by co-immunoprecipitation. This suggests that RYBP is dispensable for the assembly and activity of the Siesta-RING1 complexes *in vivo* but makes the complexes more resistant to dissociation during protein extract preparation. The increased stability may explain why RYBP-RNF2 complexes perform better in ubiquitylating H2AK119 *in vitro* (Gao *et al*., 2012; Rose *et al*., 2016).

In *Drosophila* cells, RING1 protein appears to be limiting and unstable when not incorporated into a complex. This makes distinct PCGF proteins compete for the common RING1 partner. Since PRC1 and Siesta-RING1 complexes have distinct functions, it is important to maintain the relative abundance of the two kinds of complexes in the cell. This balance appears to be regulated, at least in part, through the expression of *Psc* and *Su(z)2*. These two genes are located next to each other in a locus that includes multiple PREs bound by PRC1 and PRC2 (Park *et al*., 2012; Schwartz *et al*., 2006). This configuration generates a feedback loop where a reduction in Psc or Su(z)2 increases the transcription of the *Psc* and *Su(z)2* genes (Ali and Bender, 2004), tilting the balance towards PRC1 assembly. On the other hand, when Siesta levels decline, more RING1 protein is available for incorporation into PRC1, increasing its intranuclear concentration and enhancing transcriptional repression of *Psc* and *Su(z)2* genes. This feedback loop may be evolutionarily conserved. In humans, the *BMI1* gene, encoding one of the two PCGF proteins specific to canonical PRC1 and an ortholog of *Psc/Su(z)2,* is also regulated by the Polycomb system (Barrasa et al., 2023). Whether the second human ortholog, *MEL18*, is similarly regulated by the Polycomb system, and whether loss of PCGF proteins specific to the variant RING1/RING2 complexes reduces transcription of the *BMI1* or *MEL18* genes are fascinating open questions to address in the future.

The locomotion defect of the Siesta-KO mutants parallels behavioural abnormalities caused by mutations of the mammalian *Autism Susceptibility Gene 2* (*AUTS2*) (Liu *et al*., 2021) and locomotion defects of the *Drosophila Tay bridge* (*Tay*) (Poeck et al., 2008) genes encoding corresponding subunits of one of the Siesta-RING1 complexes (Gao *et al*., 2014; Kang *et al*., 2022). Consistent with our findings, mammalian AUTS2-PCGF3/5-RING1/2 complexes (analogues of Tay-Siesta-RING1 complex) act independently of H2AK119 ubiquitylation and appear to stimulate transcription of a subset of genes (Gao *et al*., 2014; Liu *et al*., 2021). While we have clear evidence that Siesta complexes (likely the Tay-Siesta-RING1) control larval locomotion independently of H2AK118 ubiquitylation, it is premature to conclude that their action does not require ubiquitin E3 ligase activity. It is possible that, in addition to H2AK118, the Tay-Siesta-RING1 complexes ubiquitylate yet unknown non-histone protein(s). These proteins may represent the biologically relevant target of Siesta-RING1 ubiquitylation, while H2AK118ub may be an abundant side product. It is worth pointing out that the truncated RING1-I48A protein (or the analogous truncated version of mammalian orthologues), used to benchmark the effects of E3 inactivation of RING1 complexes, shows drastically impaired E3 ligase activity towards H2AK118. However, this may not hold true for the hypothetical non-histone substrate. The same line of reasoning applies to the E3 ubiquitin ligase activity of canonical PRC1. The findings and mutants presented in our work provide an important framework to test these hypotheses and search for the non-histone targets of RING1 ubiquitylation.

## METHODS

### Generation of *RYBP* mutant alleles

The two gRNAs were synthesized with GeneArt Precision gRNA Synthesis Kit (Invitrogen). The following oligonucleotides were used to amplify the DNA templates for in vitro transcription: gRNA1-fw (5’-TAATACGACTCACTATAGGGATTGCGGTTACTCCA-3’), gRNA1-rev (5’-TTCTAGCTCTAAAACCTGGAGTAACCGCAATCC-3’), gRNA2-fw (5’-TAATACGACTCACTATAGGACAGCCGGAGTTAGAG-3’), gRNA2-rev (5’-TTCTAGCTCTAAAACCCTCTAACTCCGGCTGTC-3’). To prepare gRNA-Cas9 RNPs, a mixture of 1μg/μl TrueCut Cas9 (Invitrogen) with 0.25ng/μl of each gRNA was incubated at room temperature, and the solution cleared by centrifugation. The resulting supernatant was used for injection of *w^1118^* embryos (BDSC strain #5905). Individual F0 males were crossed with isogenic balancer stock *w^1118^; sna^Sco^/SM6a* females (BDSC strain #5907). Their male progeny was again individually crossed to *w^1118^; sna^Sco^/SM6a* females, and after 7 days of breeding, males were sacrificed for a PCR deletion screen using primers 5’-TCGAAGTAGGGTTCTCTGCC-3’ and 5’-TGAGTGCGGCTTTAATTGCGTT-3’. The homozygous mutant strains were further analysed by Sanger sequencing.

### Survival assay

To assess the viability and determine the lethal stage, first instar larvae of a genotype of interest were individually picked based on the GFP expression and put into apple juice agar plates supplemented with a small block of regular fly media. After 36-40 hours, second instar larvae were transferred to vials containing fly media (50 animals per vial) and their development followed by recording the number of prepupae, pupae and hatched adults.

### Generation of Twin-Strep-Myc-Siesta expressing transgenes

To generate a transgenic construct expressing tagged Siesta protein under control of the endogenous *Sist* promoter, the 5’ and 3’ parts of the *Sist* locus (3L:16,587,861-16,590,261, *dm6* genomic corrdinates) were PCR amplified separately using genomic DNA as a template and the L(3)73 genomic-For (5’-CGGGCCCCCCCTCGACCACTATGCACATAGACAC-3’) and iPCR-L(3)73-Rev (5’-CATGCTGGCCCCCCGATTG-3’) primers for the 5’ part and L(3)73Ah genomic-Rev (5’-CGGTGGCGGCCTCGACAAAGGCTACCACGTCCT-3’) and iPCR-L(3)73-For (5’-GAGCGGCGCGTGAAGCTGAAG-3’) for the 3’ part, respectively. The resulting fragments were combined to flank the synthetic DNA sequence encoding tandem Twin-Strep and Myc tags (amino acid sequence ASWSHPQFEKGGGSGGGSGGGSWSHPQFEQKLISEEDLEERRVK) and introduced into pW-attB vector (Savitsky et al., 2016) by In-Fusion reaction, yielding the Endo-L(3)73Ah-pW-attB construct. The construct was injected into *y1, M{vas-int.Dm}ZH-2A, w-; M{3×P3-RFP.attP}ZH-51D* embryos by BestGene.

### Generation of maternal, zygotic *siesta* mutants

The Siesta-KO^mz^ mutants were generated as described by (Gambetta and Muller, 2014). To this end, the pUMR-FLAP construct (Gambetta and Muller, 2014) was digested with EcoRI and combined with the DNA fragment produced by PCR using l(3)_FLAP_For (5’-TACGAGATCTGAATTCCACTATGCACATAGACACCATA-3’) and l(3)_FLAP_Rev (5’-GAACTTCGAAGAATTCAAAGGCTACCACGTCCTC-3’) primers and the DNA of Endo-L(3)73Ah-pW-attB as a template using In-Fusion reaction. The resulting construct was injected into *y1, M{vas-int.Dm}ZH-2A, w-; M{3×P3-RFP.attP}ZH-51D* embryos by BestGene.To generate *w^−^/w^−^; +/+; sist^1012^, {UASp-FLPo, w^+^}VK33, y^+^/TM3, w, Ser, Act::GFP, e* females; *+/+; sist^1012^/ {UASp-FLPo, w^+^}VK33, y^+^*females were crossed to *+/+; Su(z)12^4^, P{w^+^mC UAS-RAS^V12^}/TM3, Sb, e* males. *+/+; sist^1012^, {UASp-FLPo, w^+^}VK33, y^+^/ TM3, Sb, e* males were then crossed to *sist^1012^/TM3, w, Ser, Act::GFP, e* females to generate *+/+; sist^1012^, {UASp-FLPo, w^+^}VK33, y^+^/ TM3, w, Ser, Act::GFP, e* flies.

To generate *P{<sist::siesta, UAS::GFP<}ZH-51D; P{w^+mC^ = GAL4::VP16-nos.UTR}CG6325^MVD1^, sist^612^* males; *+/+; sist^612^/TM3, w, Ser, Act::GFP, e* females were crossed to *If/ CyO ; P{w^+mC^ = GAL4::VP16-nos.UTR}CG6325^MVD1^/ TM3, Sb, e* males. Then, *+/ CyO; sist^612^/ P{w^+mC^ = GAL4::VP16-nos.UTR}CG6325^MVD1^* females were crossed to *+/+; Su(z)12^4^, P{w^+^mC UAS-RAS^V12^}/TM3, Sb, e* males. The resulting *+/ CyO; sist^612^, P{w^+mC^ = GAL4::VP16-nos.UTR}CG6325^MVD1^/ TM3, Sb, e* males were crossed to *+/+;sist^1012^/TM3, w, Ser, Act::GFP, e* females to generate *+/ CyO; sist^612^, P{w^+mC^ = GAL4::VP16-nos.UTR}CG6325^MVD1^*/ *TM3, w, Ser, Act::GFP, e* females. These were then crossed to *If/ CyO ; TM6, Tb/MKRS, Sb* to generate *+/CyO; sist^612^, P{w^+mC^ = GAL4::VP16-nos.UTR}CG6325^MVD1^/ MKRS, Sb* females. Later, they were crossed to *P{<sist::siesta, UAS::GFP<}ZH-51D/If;+/ TM3, w, Ser, Act::GFP, e* males to generate *P{<sist::siesta, UAS::GFP<}ZH-51D/CyO; sist^612^, P{w^+mC^ = GAL4::VP16-nos.UTR}CG6325^MVD1^/TM3, w, Ser, Act::GFP, e.* These were then backcrossed to each other to generate the final *P{<sist::siesta, UAS::GFP<}ZH-51D; P{w^+mC^ = GAL4::VP16-nos.UTR}CG6325^MVD1^, sist^612^* flies.

To generate the Siesta-KO^mz^ embryos, the *P{<sist::siesta, UAS::GFP<}ZH-51D; P{w^+mC^ = GAL4::VP16-nos.UTR}CG6325^MVD1^, sist^612^*males were crossed to the *w^−^/w^−^; +/+; sist^1012^, {UASp-FLPo, w^+^}VK33, y^+^/TM3, w, Ser, Act::GFP, e* females. From the resulting F1 progeny, males and females of the *P{<sist::siesta, UAS::GFP<}ZH-51D/+; P{w^+mC^ = GAL4::VP16-nos.UTR}CG6325^MVD1^, sist^612^/sist^1012^, {UASp-FLPo, w^+^}VK33, y^+^* genotype were crossed with each other, which resulted in embryos that lack maternal and zygotic Siesta protein. Control embryos were generated by crossing Oregon R females with *P{<sist::siesta, UAS::GFP<}ZH-51D; P{w^+mC^ = GAL4::VP16-nos.UTR}CG6325^MVD1^, sist^612^* males. The F1 progeny were then crossed with each other. Since there was no FLP recombinase source, the transgenic cassette was not excised (Figure S7).

### Locomotion assay

The locomotion of *Drosophila* second instar larvae was recorded and analysed as previously described (Brooks et al., 2016). Briefly, second instar larvae were transferred to apple juice agar plates supplemented with 0.1% bromophenol blue and allowed to acclimate for 30-60 sec. A mobile phone with a 48-megapixel camera (480 x 640 pixels recording resolution) was used to capture videos of moving larvae. Four to six larvae were tracked in every video for a minimum of 90 seconds in a thermally controlled room set to 25 C°. The WrMTrck ImageJ plugin (Nussbaum-Krammer et al., 2015) was used to analyse the videos. For each larva, the trimmed mean motion speed was calculated by omitting video frames with 10% of the highest and 10% of the lowest speeds to reduce background noise. The locomotion of first instar larvae was assayed as follows. The first instar larvae were placed on apple juice agar plates and allowed to acclimate for 30-60 seconds. Videos were acquired using a FLIR camera (480 x 640 pixels recording resolution) mounted on a Zeiss Stemi-508 microscope as described (Pu et al., 2023) and analysed using WrMTrck ImageJ plugin as described above except that all video frames were analysed and mean motion speed was calculated. To examine the hypoxia-evoked locomotory response, the larvae were sealed in a microfluidic chamber, and defined gases were delivered as described (Pu *et al*., 2023).

### Immunostaining and microscopy

*Drosophila* embryos were fixed and stained as described (Dorafshan et al., 2019b) with the following modifications. The embryos were dechorionated for 3 min in 7% sodium hypochlorite, and the blocking solution contained 5% Newborn Calf Serum in PBST (137mM NaCl, 2.7mM KCl, 10mM Na_2_HPO_4_, 2mM KH_2_PO_4_, 0.1% Tween-20) instead of normal goat serum. The stained embryos were mounted on a glass slide and imaged with a Leica SP8 confocal microscope. Immunostaining and imaging of polytene chromosomes were done as described in (Dorafshan et al., 2019a). The antibodies used are listed in Table S2.

### Generation of Siesta-KO, RING1-KO and control cell lines

The Siesta-KO cell lines were derived from homozygous *sist^612^*embryos using the Ras^V12^-mediated transformation approach by Simcox and colleagues (Simcox *et al*., 2008) with modifications described in (Kahn *et al*., 2016). Ras^V12^-transformed but otherwise wild-type control cell line (Ras17) was derived in parallel using the same procedure.

Cultured cell lines homozygous for the deletion of the *Sce* gene (RING1-KO cells) were derived by CRISPR/Cas9-mediated genome editing of Ras17 cells. Generation of *Drosophila* knock-out cell lines using CRISPR/Cas9-mediated genome editing has two challenges. First, cultured *Drosophila* cells are difficult to clone due to poor cell proliferation in dilute cultures. Second, the frequency of generating deletions in both copies of a gene is low. This is likely due to the strong somatic pairing of homologous chromosomes, which causes most of the double-strand DNA breaks induced by Cas9 to be repaired by homology-directed DNA repair using the unedited homologue as a template. To overcome these limitations, we developed a new cell cloning protocol and a two-step CRISPR-Cas9 editing strategy, which is described below.

To isolate single-cell clones, cultured *Drosophila* cells are diluted 100, 1000 and 10000 times and plated in a 6-well plate covered with 0.1% gelatine. After 5-7 days, clusters of cells originated from mitotic divisions of individual cells are manually picked under a dissection microscope and reseeded in a 96-well plate. In 2-3 weeks, successfully proliferating clonal cultures are reseeded into a 24-well plate and genotyped. For the two-step CRISPR/Cas9 editing, four single guide RNAs are designed to generate a deletion of interest (Figure S12). In the first editing round, the cells are transfected with the Cas9/gRNA complex that includes an outer pair of gRNAs (Figure S12) and the resulting cell population is used to derive single-cell clonal cultures. At this stage, all of the resulting cultures are heterozygous with only one allele bearing the desired deletion. In the second step, two independently isolated heterozygous clones are treated with the inner pair of sgRNAs. At this step, the wild-type template for homology-directed DNA repair is not available, so in most cases, the deletion allele obtained in the first step is used as a template instead. The resulting cell population is used to derive single-cell clonal cultures. After the second round of editing and cloning, more than half of the derived cultured cell lines are homozygous for the desired deletion.

To generate cultured cell lines homozygous for the deletion of the *Sce* gene (RING1-KO cells), Ras17 cells were subjected to two rounds of CRISPR/Cas9-mediated genome editing as outlined above. For each editing round, 50 picomoles of each single guide RNA (Synthego), were mixed with 60 picomoles of Cas9 nuclease (IDT) and delivered to one million cells by electroporation with Neon Transfection System 10µl kit from ThermoFisher Scientific with 1800V pulse for 10 milliseconds twice according to manufacturer instructions. In the first editing round, the Cas9 complexes with gRNA-dSce-ex1.1 (5’-UGGUGUGAAAAUGACGUCGC-3’) and gRNA-dSce-ex2.1 (5’-CUAUGGAAAUGUAUUACUCG-3’) were used for treatment, which resulted in the *Sce* deletion from one of the homologous chromosomes, detected in several clonal cell cultures (Figure S13). Two such cultures (cell lines Sce-KO63-2 and Sce-KO66) were used for the second editing round using Cas9 in the complex with gRNA-dSce-ex1.2 (5’-CCCGGCGCCAAACAAAACGU-3’) and gRNA-dSce-ex2.2 (5’-AAAGCAUACAGACAUUAGAA-3’). The resulting cell population was used to derive single-cell clonal cultures, some of which were genotyped by PCR amplification of the junction and sequencing the PCR fragments (Figure S13). Half of the genotyped clonal cultures were homozygous for the deletion allele obtained in the first editing round due to homology-directed repair of the DNA breaks produced during the second editing round. The other half of the clones harboured two different alleles, one allele with the original deletion from the first editing round and the second allele originated from non-homologous end joining of dsDNA breaks produced during the second editing round. The nucleotide sequences of PCR primers are listed in Table S3. All cell lines were cultured at 25°C in Schneider’s medium (BioConcept) supplemented with 10% of heat-inactivated foetal bovine serum (Sigma), streptomycin (0.1mg/ml), and penicillin (100 units/ml) (Gibco) under sterile conditions.

### Protein co-immunoprecipitation

#### Embryo collection

Embryos were collected twice daily from 20 ventilated 250ml Erlenmeyer flasks assembled with apple juice plates with a knob of yeast paste and populated with three to four hundred 3-5 day old flies. The harvested embryos were placed in a 70 µM Corning cell strainer, washed with running deionised water, rinsed with Embryo Wash Solution (0.04% Triton X-100, 120mM NaCl) and again with running deionised water. Then embryos were partially dried and packed into a “tablet” by blotting the mesh with paper tissues, weighed, wrapped in aluminium foil and flash frozen in liquid nitrogen. Frozen embryos were stored at -80^0^ C until protein extraction.

#### Nuclear protein extraction

Approximately 12g of embryos were placed in a DIY nylon mesh-bottomed basket and submerged in a glass with 3.5 % bleach for 4 min, while breaking lumps and gently stirring with a plastic spoon, to remove the chorions. The dechorionated embryos were washed extensively with deionized water and transferred into pre-chilled 40 ml Wheaton glass homogenizer. Embryos were resuspended in 10 ml of Embryo buffer (15 mM HEPES pH 7.6, 10 mM KCl, 5 mM MgCl_2_, 0.1 mM EDTA, 0.5 mM EGTA, 350 mM Sucrose, 1 mM DTT, 0.5 mM PMSF, 1x protease inhibitor cocktail cOmplete, EDTA-free (Roche) and homogenized with 10 strokes of pestle A, followed by 20 strokes of pestle B. The resulting homogenate was filtered through 70µm cell strainer, debris re-suspended in 10ml of embryo buffer, and given an additional 10 strokes with pestle B, followed by filtration. The combined homogenate was centrifuged for 15min at 10 000g and 4 °C. The supernatant was carefully aspirated and discarded, and fat deposits were wiped from the walls with tissue paper. Nuclei were resuspended in 10ml of Embryo buffer, trying not to disturb the yellow egg yolk layer at the bottom, and the centrifugation was repeated. The resulting nuclei were resuspended in 2 ml of Low-salt Nuclear Buffer (15 mM HEPES, pH 7.6, 20% glycerol, 1.5mM MgCl_2_, 20 mM KCl, 0.1 mM EDTA, 1 mM DTT, 0.5 mM PMSF, complete protease inhibitor cocktail, EDTA-free (Roche)) and the concentration of KCl in the suspension adjusted to 400 mM with 3M KCl solution. The suspension was incubated on ice with occasional gentle stirring (every 5-10 minutes) and then another 30 minutes inside a pre-chilled ultra-centrifuge (meanwhile, a vacuum was reached). The suspension was centrifuged for 1 hour at 4 °C, 100 000g to separate into four layers. Of those, the dense top fat layer was mechanically removed with a 5µL microbiological loop, the transparent off-white nuclear extract (about 2.2 - 2.5 ml total) was split into 500µL aliquots, flash frozen in liquid nitrogen and stored at -80^0^ C, while opaque greyish liquid phase and solid bottom phase were discarded.

#### Antibody crosslinking to magnetic beads

The crosslinking was performed according to the manufacturer’s instructions with the following modifications. Dynabeads (Thermo Fisher) were separated from the storage solution by incubation on a magnetic stand and washed 2 times in 200µl of Wash Buffer (1xPBS, 0.02% NP40, 1 mM DTT, 0.5 mM PMSF, complete protease inhibitor cocktail, EDTA-free (Roche)), and resuspended in 200µl of Wash Buffer. 1µg of anti-RING1 antibody or 1µg of normal rabbit IgG (Sigma-Aldrich NI01-100UG) was added to the bead suspension and incubated for 30 min at room temperature while slowly rotating. Antibody-bound Dynabeads were washed one time with Wash Buffer, 2 times with 200µl Conjugation Buffer (20 mM sodium phosphate, 0.15M NaCl, pH 8), and resuspended in 125 µl Conjugation Buffer. The crosslinking was initiated by adding 125µl of 10mM BS3 (bis(sulfosuccinimidyl)suberate) solution (Thermo Fisher), followed by slow rotation at room temperature for 30 minutes. The reaction was stopped by quenching with 12.5µl of 1M Tris, pH 7.5, for 15 min with slow rotation. The resulting affinity resins were washed 3 times with Low-salt Nuclear Buffer supplemented with 0.02% NP40 and 1 mM DTT and stored at 4 °C.

#### Immunoprecipitation

Nuclear extracts were defrosted on ice and dialysed 2 times for 2-3 hours against 14 ml of 25 mM HEPES-KOH, pH 7.6; 0.1 M KCl; 1.5 mM MgCl_2_; 0.1 mM EDTA; 20% (v/v) glycerol, 1 mM DTT, 0.5 mM PMSF, cOmplete protease inhibitor cocktail, EDTA-free (Roche) using Slide-A-Lyzer™ MINI Dialysis Devices, 10K MWCO (Thermo Fisher) placed on ice with slow rocking. The affinity resins were washed 2 times with Low-salt Nuclear Buffer supplemented with 0.02% NP40 and 1 mM DTT and combined with 100μl of the dialysed extracts pre-cleared by centrifugation. The resulting suspensions were adjusted to contain 0.02% of NP40 and incubated overnight with slow rotation at 4 °C. The resins were washed 3 times with 500µl of Wash Buffer and once with Wash Buffer diluted 1:10 with water. For washing, Dynabeads were gently resuspended in the buffer by pipetting and separated on a magnetic stand, while Strep-Tactin Sepharose (IBA) beads were rotated for 5 minutes before centrifugation (2 min, 500g). The protein bound to antibody-coupled Dynabeads was eluted with 100µL of 1x NuPAGE LDS sample buffer (Invitrogen) supplemented with 5% β-mercaptoethanol at 98^0^C for 10 min. The protein bound to Strep-Tactin Sepharose was eluted by 10 min incubation with 40µL of 2 x NuPAGE LDS sample buffer followed by incubation with an additional 50µL 1 x NuPAGE LDS sample buffer, and combining the eluates. To prepare the matching immunoprecipitation input samples, the dialysed nuclear extracts were combined with 4 volumes of acetone (pre-chilled to -20°C) and subjected to protein precipitation overnight at -20°C. The precipitated protein was pelleted by centrifugation, washed with 50% acetone and air-dried for 10 minutes. Protein pellets were resuspended in 1x NuPAGE LDS sample buffer (200µl per 1 input volume) for 10 min at 98 °C, homogenised with a pestle, vortexed, heated for an additional 10 min, and centrifuged to remove the remaining insoluble material.

### qRT–PCR

Total RNA from 5×10^6^ cells was isolated using Trizol (Invitrogen), and 2µg was used for random primed synthesis of cDNA with First Strand Synthesis Kit (ThermoFischer). The control reaction, omitting reverse transcriptase, was always run in parallel. cDNA was treated with RNase A to remove the RNA template, purified with the Zymo PCR purification kit and eluted with 200µl of elution buffer. The amount of cDNA for a gene of interest in the given preparation was analysed by qPCR and was expressed as a fraction of RpL32 cDNA. Serial dilutions of genomic DNA were used to make the standard curve. qPCR analysis was performed using a BioRad CFX Connect Real Time PCR instrument in a total reaction volume of 10µl containing 4µl of cDNA solution, 1x qPCRBIO SYGreen Master Mix (PCRBioSystems), and 200nM of corresponding primers (sequences indicated in Table S3).

### ChIP

#### Preparation of MNase-digested chromatin from crosslinked Drosophila cultured cells

Cells were crosslinked by adding the formaldehyde solution directly to the cell culture to a final concentration of 1.8% and incubating for 10 minutes at 25^0^C. The reaction was stopped by adding glycine pH 7.0 to a final concentration of 0.125M. Crosslinked cells were washed once in cold ChIP Wash Buffer A (10mM Hepes pH7.0, 10mM EDTA pH8.0, 0.5mM EGTA pH8.0, 0.25% Triton X-100) for 10 minutes at 4^0^C and once in cold ChIP Wash Buffer B (10mM Hepes pH7.6, 100mM NaCl, 1mM EDTA pH8.0, 0.5mM EGTA pH8.0, 0.01% Triton X-100) for 10 minutes at 4^0^C. Approximately 0.5 ml of pelleted crosslinked cells were resuspended in 5 ml cold TE buffer (10mM Tris-HCl pH8.0, 1mM EDTA pH8.0) and incubated with 0.1u/ml MNase (Sigma N3755, MNase powder was reconstituted with water and adjusted to 50% glycerol to achieve 0.1u/µl MNase solution and stored at -20^0^C) and 2mM CaCl_2_ at 37^0^C for 25 minutes while shaking at 1000 rpm. The reaction was stopped by adding EGTA to a final concentration of 10mM. The cells were pelleted at 1000g for 2 minutes at 4^0^C, washed with 5ml of cold TE, pelleted again and resuspended in 4ml of cold TE. The cells were lysed by ultrasound using Branson sonicator (9 cycles of 20 seconds ON followed by 40 seconds OFF with 40% amplitude in ethanol-ice bath). The cell lysates were adjusted to 5ml RIPA buffer by sequential addition of Triton X-100 to 1%, DOC to 0.1%, NaCl to 140mM and SDS to 0.1%, TE to 1x, incubated at 4^0^C for 10 minutes on a rotating wheel and centrifuged for 5 minutes at 12000xg to remove insoluble residues. Soluble cell lysates (crosslinked chromatins) were aliquoted, frozen in liquid nitrogen and stored at -80^0^C. 100µl aliquot of each cell lysate was used to purify the DNA and estimate the DNA content.

#### Preparation of MNase-digested chromatin from crosslinked Drosophila embryos

Approximately 0.7g of 16-18 hour old *Drosophila* embryos were dechorionized by incubation in 3-2.5% Na-hypochlorite solution for 3 minutes at room temperature, washed twice with 0.4% NaCl, 0.03% Triton X-100, and crosslinked with 1.8% formaldehyde solution in 50 mM Hepes pH7.6, 1mM EDTA pH8.0, 0.5mM EGTA pH8.0, 100mM NaCl by shaking for 20 minutes at room temperature in the presence of n-heptane. The reaction was stopped by adding glycine pH7.0 to a final concentration of 0.125M. Crosslinked embryos were washed once in cold ChIP Wash Buffer A for 10 minutes at 4^0^C, and once in cold ChIP Wash Buffer B for 10 minutes at 4^0^C. The crosslinked embryos were resuspended in 5ml of cold ChIP Wash Buffer B, transferred into 7ml homogeniser, and disrupted by several strokes of a tight pestle. The crosslinked embryonic cells were pelleted at 2000g for 2 minutes at 4^0^C, washed once with 5 ml of cold TE buffer, resuspended in 3.5ml of cold TE buffer, and incubated at 37^0^C with 0.1u/ml MNase and 2mM CaCl_2_ for 25 minutes while shaking at 1000rpm. The reaction was stopped by adding EGTA to a final concentration of 10mM. The crosslinked embryonic cells were precipitated by centrifugation at 1000g for 2 minutes at 4^0^C, washed with 5 ml of cold TE, precipitated again and resuspended in 3ml of cold TE. The cells were lysed by ultrasound using Branson sonicator (9 cycles of 20 seconds ON followed by 40 seconds OFF with 30% amplitude in ethanol-ice bath). The resulting lysates were adjusted to 3.5ml of RIPA by sequential addition of Triton X-100 to 1%, DOC to 0.1%, NaCl to 140 mM and SDS to 0.1%, TE to 1x and processed as described above for the crosslinked cultured cell lysates.

#### Immunoprecipitation

ChIP was done essentially as described in (Schwartz *et al*., 2006) with the following modifications. A standard amount of crosslinked cell lysates containing either 100µg of DNA (for cultured cells) or 50µg of DNA(for embryos) was used for immunoprecipitations in a total volume of 500µl of RIPA buffer (140mM NaCl, 10mM Tris-HCl pH8.0, 1mM EDTA, 1% Triton X-100, 0.1% SDS, 0.1% DOC). The resulting immunoprecipitated DNA was dissolved in either 400µl of water for qPCR analysis or in 40µl of water for ChIP-seq library preparation. The nucleotide sequences of primers used for qPCR analysis are listed in Table S3. Serial dilutions of DNA isolated from crosslinked cell lysates (input) were used to make the standard curves, and the abundance of specific regions in the immunoprecipitated DNA was expressed as a fraction of input. Rabbit polyclonal antisera against RING1 and Psc were raised by immunisation with recombinant GST fusion proteins containing amino acid residues 150-280 of RING1 and amino acid residues 821-1021 of Psc, respectively. The antibodies were further affinity purified as described in (Poux et al., 2001) and are available for purchase from Agrisera AB. All other antibodies were purchased from vendors listed in Table S2.

### ChIP-seq

4 ng of ChIP DNA and corresponding Input DNA were used to prepare sequencing libraries. The library list and the corresponding fractions of the total material used for preparation are indicated in Table S4. The libraries were prepared using NEBNext Ultra II FS DNA Library Prep with Beads (New England Biolabs #E6177S) according to the manufacturer’s instructions, with minor modifications described below. To get the libraries of fragments with an average size of approximately 350-400bp, the DNA samples were fragmented for 10 minutes at 37^0^C using NEBNext Ultra II FS Enzyme Mix from the kit. The Illumina adaptor was diluted 25-fold before the ligation to the DNA fragments. The libraries were multiplexed using NEBNext Multiplex Oligos for Illumina, Set 1 (New England Biolabs #E7335S) and Set 2 (New England Biolabs #E7500S) during 9 cycles of amplification and cleaned up twice with the supplied magnetic beads to remove the primers and adaptor-dimers. A fraction of each library was used to prepare 10µl of 10nM solution. All individual 10nM libraries were combined proportionally to one 10nM library pool. Fractions of each ChIP DNA and Input DNA used to prepare 10nM library are listed in Table S4. The pool was sequenced on 1 lane of 10B flow cell using the NovaSeq X Plus system and XLEAP-SBS sequencing chemistry (Illumina).

### ChIP-seq data analysis

#### Read alignment

Sequencing reads were aligned to the *Drosophila melanogaster dm6* reference genome using bowtie2 (v2.4.4) (Langmead and Salzberg, 2012) with arguments set to *–phred33 --no-discordant --very-sensitive-local -p 32*. The alignment output was further filtered using samtools (v1.3.1) to remove reads with a mapping quality score less than 30 *(-h -b -@ 32 -q 30*). The resulting filtered BAM files were used as input to generate bedGraph profiles using the MACS3 (Zhang et al., 2008) pileup command with the BAMPE output option. Aligned reads were visualised with IGB (v.10.1.0) (Freese et al., 2016). Distribution of ChIP-seq signals at the whole chromosome scale was visualised with the R package chromoMap v0.3 (Anand and Rodriguez Lopez, 2022).

#### ChIP-seq signal normalization and filtering

For quantitative comparison of ChIP-seq signals for the same antibody from cells with different genetic backgrounds, we adopted the “sans-spike-in” approach for quantitative ChIP-sequencing (siQ-ChIP) proposed by Dickson and colleagues (Dickson *et al*., 2023; Dickson *et al*., 2020). siQ-ChIP aims to estimate ChIP yields (*Y*) for every genomic position. It postulates that ChIP yield *Y(i)* for the genomic position *i* corresponds to the number of DNA fragments recovered in a ChIP reaction that overlap this position *R_IP_(i)*.

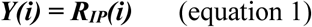

*R_IP_(i)* could be determined by sequencing all DNA molecules recovered in a ChIP reaction. In practice, it is customary to sequence just a fraction of all DNA immunoprecipitated in one reaction, and often only the nucleotide sequences of the ends of the immunoprecipitated DNA fragments are determined. Therefore, typically, the sequencing assay provides the information on the number of sequence tags *R’_IP_ (i)* that overlap position *i* rather than *R_IP_(i)*. *R_IP_(i)* is proportional to *R’_IP_(i),* and the two values are connected by the scaling coefficient *S_IP_*. This coefficient depends on several factors, for example, sequencing depth, the fraction of the library taken for sequencing, and the fraction of the ChIP DNA used to prepare the library. Considering the above, we can rewrite equation 1 as follows:

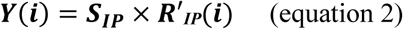

We posit that, under the standard conditions (i.e. all ChIP reactions were performed with the same amount of antibody and chromatin lysate, the immunoprecipitated DNA was uniformly fragmented before library construction, all sequencing libraries adjusted to the same concentration (10nM) and sequenced in parallel as part of the same pool), *S_IP_* is the same for all genomic positions and can be calculated as follows:

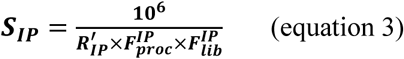

where *R’_IP_* is the total number of sequence tags obtained from sequencing of the corresponding library, *F_proc_* is a fraction of the ChIP DNA used to prepare the sequencing library, *F_lib_* is a fraction of the sequencing library used to prepare 10nM library solution, and the numerator 10^6^ to scales the yields to one million of sequenced reads. Combining equations 2 and 3 allows us to express the estimated ChIP yields *Y(i)* for any genomic position *i* as a function of variables directly measured in a the assay:

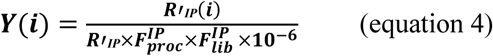

Thus, for each ChIP sample, the *F^IP^_proc_*, *F^IP^_lib_* values were recorded during library preparation, and the total number of sequenced reads *R’_IP_* was reported after sequencing. The corresponding values are indicated in Table S4. The *R’_IP_(i)* values were calculated using bedtools (v2.30.0) https://bedtools.readthedocs.i<u>°</u> using a two-step procedure. First, the genome was segmented into 50bp bins using the *bedtools makewindows -w 50* command. Second, the number of sequence tags for each bin *i* was calculated using the *bedtools map -o mean* command.

For consistency, the read distributions obtained after sequencing DNA from the corresponding chromatin input materials were scaled as described above. Although in this case, the resulting values have no biological meaning. For all the downstream analyses and visualisation, the normalized ChIP-seq signals (i.e. genomic distributions of estimated ChIP yields) were filtered to remove spurious peaks corresponding to highly repeated genomic regions. To this end, we removed all 50bp bins whose scaled read counts in the corresponding chromatin input data set exceeded the mean plus three standard deviations for this data set.

#### Definition of bound regions

Regions significantly enriched by ChIP with anti-Psc and anti-RING1 antibody were identified separately for each replicate experiment using the MACS3 *callpeak* command with the following parameters: *-f BAMPE -g dm --qvalue 0.1*. Only regions with fold enrichment (FC) ≥ 3 were used for further analyses. Regions significantly enriched by ChIP with anti-H3K27me3 antibodies were defined using Epic2 software (Stovner and Saetrom, 2019) with the genome parameter set to *--genome dm6*. Similarly, only regions with fold enrichment (FC) ≥ 3 were used for further analyses. PREs were defined as regions significantly enriched by anti-RING1, anti-Psc, and anti-H3K27me3 antibodies using *bedtools intersect* function. For all H2AK118ub ChIP-seq datasets, 1kb windows with normalized ChIP-seq signals exceeding three standard deviations from the mean normalized signal of ChIP-seq assays with chromatin from RING1-KO cells were considered significantly enriched.

#### Genome segmentation by k-means clustering

The genome was divided into 1kb windows using *bedtools makewindows -w 1000* and each window was assigned a mean normalized ChIP-seq signal value computed for H2AK118ub signals from ChIP-seq assays with chromatins from the control, Psc/Su(z)2-KO and Siesta-KO cells. The mean normalized H2AK118ub ChIP-seq signal values for each genetic background were scaled using the *scale* function to standardise the data across samples. These scaled values were then used to segregate the 1kb windows into 3 groups by unsupervised k-means clustering using the *stats* package (v.4.4.3) https://www.R-project.org/ and the following parameters: *centers = 3 and nstart = 25*. The 1kb windows with associated ChIP-seq signal values and cluster assignments are listed in Supplementary file 1.

## TRIP

### Cell transfection

The control (Ras17), Psc/Su(z)2-KO, Siesta-KO, and dRING-KO cells were cultured at 25°C in Schneider’s medium (BioConcept) supplemented with 10% of heat-inactivated foetal bovine serum (Sigma), streptomycin (0.1mg/ml), and penicillin (100 units/ml) (Gibco) under sterile conditions. For transfection, 2 × 10^6^ cells were seeded in a 6-well plate one day before the procedure. Transfection was performed using X-tremeGENE™ HP DNA Transfection Reagent (Sigma) according to the manufacturer’s instructions. Cells of one well were transfected with 2μg of the equimolar DNA mixture of the pPB-MtnA-eGFP-PI-14-BC-libr, pPB-MtnA-eGFP-PI-16-BC-libr, pPB-Hsp70-eGFP-PI-11-BC-libr and pPB-Hsp70-eGFP-PI-3-BC-libr (Supplementary files 2-5) barcoded plasmid libraries, and 0.4μg of the DNA of the construct to express the piggyBac transposase fused to histone H1 under control of the *Hsp70Bb* promoter (phsp70-pBac-H1, Supplementary file 6). Cultures of cells transfected with mixtures lacking the construct expressing the piggyBac transposase (mock-transfected) and non-transfected cells were grown in parallel. 24 to 48 hours after transfection, the treated and control cell cultures were subjected to a heat shock at 37°C for 2 hours to induce the transposase expression. The cell cultures were allowed to grow for 3-4 days, after which several aliquots of approximately 10000 cells were sub-cultured to establish TRIP cell pools, each containing a unique collection of integrated transgenes. Each TRIP cell pool was further cultured for a minimum of one month to eliminate the transfected DNA that did not integrate into the genome.

### GFP induction with CuSO_4_

To induce transcription of the transgenic *GFP*, 3×10^6^ cells plated in triplicate two days earlier were treated with 0.5mM CuSO_4_ for 2 hours at 25^0^C. The control cell cultures were grown in the same way, but without the addition of supplementary CuSO_4_.

### RNA isolation and cDNA synthesis

For each replicate preparation, 1×10^7^ cells were collected by centrifugation and lysed in 1 ml Trizol reagent (Invitrogen, #15596026). The RNA was isolated according to the manufacturer’s instructions. Two micrograms of the RNA were used for cDNA synthesis with oligo(dT)18 primer by RevertAid H Minus First Strand cDNA Synthesis Kit (Thermofisher Scientific, # K1631). The cDNA was treated with RNase A to remove the RNA template, cleaned up with Zymo DNA purification kit (#D4034) and eluted with 30 µl of 0.1x TE.

### TRIP-mapping library preparation

Two micrograms of genomic DNA from each cell line were digested with MboI at 37^0^C for 6 hours, followed by incubation at 65^0^C to inactivate the enzyme. Digested DNA (one third of the reaction volume, 0.7 μg) was circularized in 400µl reaction with 20 units of T4 DNA ligase (in the presence of 1mM ATP, 10mM MgCl2, 1mM DTT, 60mM Tris-HCl pH 7.4) at +4^0^C for 16 hours followed by incubation at 65^0^C for 10 minutes to inactivate the ligase. The resulting circular DNA was purified by phenol-chloroform extraction and dissolved in 30µl of water. 5µl of purified circular DNA solution was used for the first round of inverse PCR to amplify the parts that included the TRIP barcodes and adjacent genomic DNA from insertion sites. 1µl of the resulting PCR product was used in the second PCR round to introduce sequencing indexes. 1µl of the resulting product from the second PCR round was used in the third round of PCR to incorporate Illumina sequencing adaptors. The list of primers and amplification conditions is indicated in Table S5. The resulting TRIP-mapping libraries were cleaned up with SPRIselect magnetic beads (Beckman Coulter) twice, adjusted to 10nM and pooled proportionally into 10nM TRIP-mapping pool. All sequenced TRIP-mapping libraries are listed in Table S6.

### TRIP-normalization library and TRIP-cDNA library preparation

2.5×10^7^ cells were lysed overnight in 1x TE with 0.5 mg/ml Proteinase K and 0.5% SDS, followed by phenol-chloroform extraction and ethanol precipitation. The resulting genomic DNA was dissolved in 1x TE (10mM Tris-HCl, pH 8.0; 1mM EDTA) and treated with 0.1 mg/ml RNase A at 37^0^C for 30 minutes. RNase A was removed by phenol-chloroform extraction, the DNA was ethanol precipitated and dissolved in 1x TE.

100ng of the purified genomic DNA or 10% of the cDNA product from the synthesis reaction (described above) were used for normalization and cDNA library preparation, respectively. The DNA was subjected to two rounds of PCR to introduce sequencing indexes and Illumina adaptors for sequencing. PCR products were cleaned up with SPRIselect magnetic beads after every PCR round and eluted with 25µl of 0.1x TE buffer. 5µl of the eluted DNA after the first PCR round were used in the second round. The libraries were adjusted to 10nM and mixed proportionally into TRIP-normalization and TRIP-cDNA pools. The libraries are listed in Table S6. The primers and amplification conditions are described in Table S7.

Fractions of TRIP-mapping, TRIP-normalization and TRIP-cDNA libraries were combined into a single 10nM pool at ratios 1:2:2. The pool was sequenced on 1 lane of 10B flow cell using the NovaSeq X Plus system and XLEAP-SBS sequencing chemistry (Illumina).

#### TRIP data analyses

As illustrated by Figure 6, the TRIP assay yields three types of sequencing reads. Although all libraries were sequenced as paired end, for the “expression” and “normalization” datasets, only single-end reverse 150bp reads were used. These reads, which contained the plasmid region and a unique barcode, were used to count the barcode frequencies in genomic DNA and cDNA samples, respectively. The third dataset contains “mapping” reads. These are paired-end 150bp reads obtained by sequencing of inverse PCR products. The “mapping” reads include the 5’ end of the piggyBac transposon and a neighbouring genomic DNA sequence in the forward read, as well as a barcode and a genomic sequence adjacent to the DpnII site in the reverse read.

The initial fastq file with reads obtained after sequencing the TRIP library pool was parsed according to the sequencing index (Table S6) using *sabre* software (https://anaconda.org/bioconda/sabre) and then by the promoter index using *awk*-based *bash* script (Supplementary file 7). The parsed fastq files with “normalization”, “expression”, and “mapping” reads were grouped by individual TRIP experiment and processed with *TRIP Analysis Software Kit (TASK)* software (Akhtar et al., 2014) with parameter and read structure settings described in Supplementary file 8. The example command line instructions to process a dataset with TASK software are provided in Supplementary file 9.

The barcode lists returned by *TASK* were filtered to remove the instances that may have arisen due to nucleotide substitutions during the PCR steps of library preparation. To this end, the following algorithm was used. First, the barcode instances were sorted according to their counts in the “normalization” sample. Second, the most frequent barcodes were ranked the highest, and for each highest-ranked barcode instance, “mutant” versions, defined as barcodes with distinct nucleotide sequences appearing at the same position (pos_r), were identified and removed. Third, the resulting lists were filtered to remove all barcode instances with a mapping quality score less than 10 (mapq_r > 10). Finally, the identical barcode instances associated with the same genomic position in two biologically independent replicate experiments were deemed to represent genuine transgenic insertions.

Each genuine transgenic insertion was assigned the TRIP signal calculated as follows:

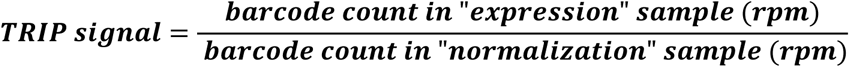

The list of genuine transgenic insertions with associated nucleotide sequence, genomic position, TRIP signal, etc, is reported in Supplementary file 10.

#### Statistical analysis and plotting

All statistical analyses were performed using R https://www.R-project.org/, and plots were generated using ggplot2 (Wickham, 2016).

## DATA AVAILABILITY

All data needed to evaluate the conclusions in the paper are presented in the paper and/or Supplementary Materials. ChIP-seq and TRIP datasets generated in this study are available from the Gene Expression Omnibus (http://www.ncbi.nlm.nih.gov/geo/), accession numbers GSE302678, GSE302873.

## ACKNOWLEDGEMENTS

We thank Alessandro Gozzo for help in the generation of *Sist^mz^*animals. We are grateful to Jürg Müller, Maria Gambetta and Mikhail Savitsky for sharing recombinant DNA constructs and fly stocks. We also thank Alexey Pindyurin and Lyubov Yarynich for sharing unpublished DNA constructs and helping with the setup of the TRIP assay. We are grateful to Mikael Lindberg (Protein Expertise Platform, Umeå University) for the production of recombinant Gst-RING1 and Gst-Psc and to Changchun Chen and Lars Nilsson for helping with locomotion tracking. This work was supported by grants from Cancerfonden (22 2285Pj to YBS and 23 2762Pj to JL), Swedish Research Council (2021-04435 to YBS and 2024-03913 to JL), Kempestiftelserna (JCK22-0055) and Umeå University Insamlingsstiftelsen to YBS. Sequencing was performed by the SNP&SEQ Technology Platform in Uppsala. The facility is part of the National Genomics Infrastructure (NGI) Sweden and Science for Life Laboratory. The SNP&SEQ Platform is also supported by the Swedish Research Council and the Knut and Alice Wallenberg Foundation. Stocks obtained from the Bloomington Drosophila Stock Center (NIH P40OD018537) were used in this study. The monoclonal antibodies against Abd-B (1A2E9, donated by S. Celniker, Lawrence Berkeley National Lab), Antp (Antp 4C3, donated by D. Brower, University of Arizona) and Prospero (MR1A, donated by C.Q. Doe, University of Oregon) were obtained from the Developmental Studies Hybridoma Bank, created by the NICHD of the NIH and maintained at The University of Iowa, Department of Biology, Iowa City, IA 52242.

## AUTHOR CONTRIBUTIONS

TGK derived RING1-KO cell lines, performed RT-qPCR, ChIP, ChIP-seq, TRIP and Western blot analysis of bulk protein levels in cultured cell lines. AnG performed all genetic analyses and Western blot analyses of H2AK118ub in second instar larvae, characterised locomotion phenotypes of Siesta mutants with help from SS and assayed the expression of homeotic genes by immunostaining. AY prepared TRIP cell pools and performed computational analyses of ChIP-seq and TRIP data. MK derived RYBP mutants and performed protein co-immunoprecipitation experiments. AlG derived Siesta-KO and control cultured cell lines, generated transgenic fly strains complementing *sist* mutations and performed polytene chromosome immunostainings. JL acquired funding and supervised MK. YBS conceived and supervised the project, acquired funding, and wrote the manuscript with input from all authors.

---

**Figure S1.**
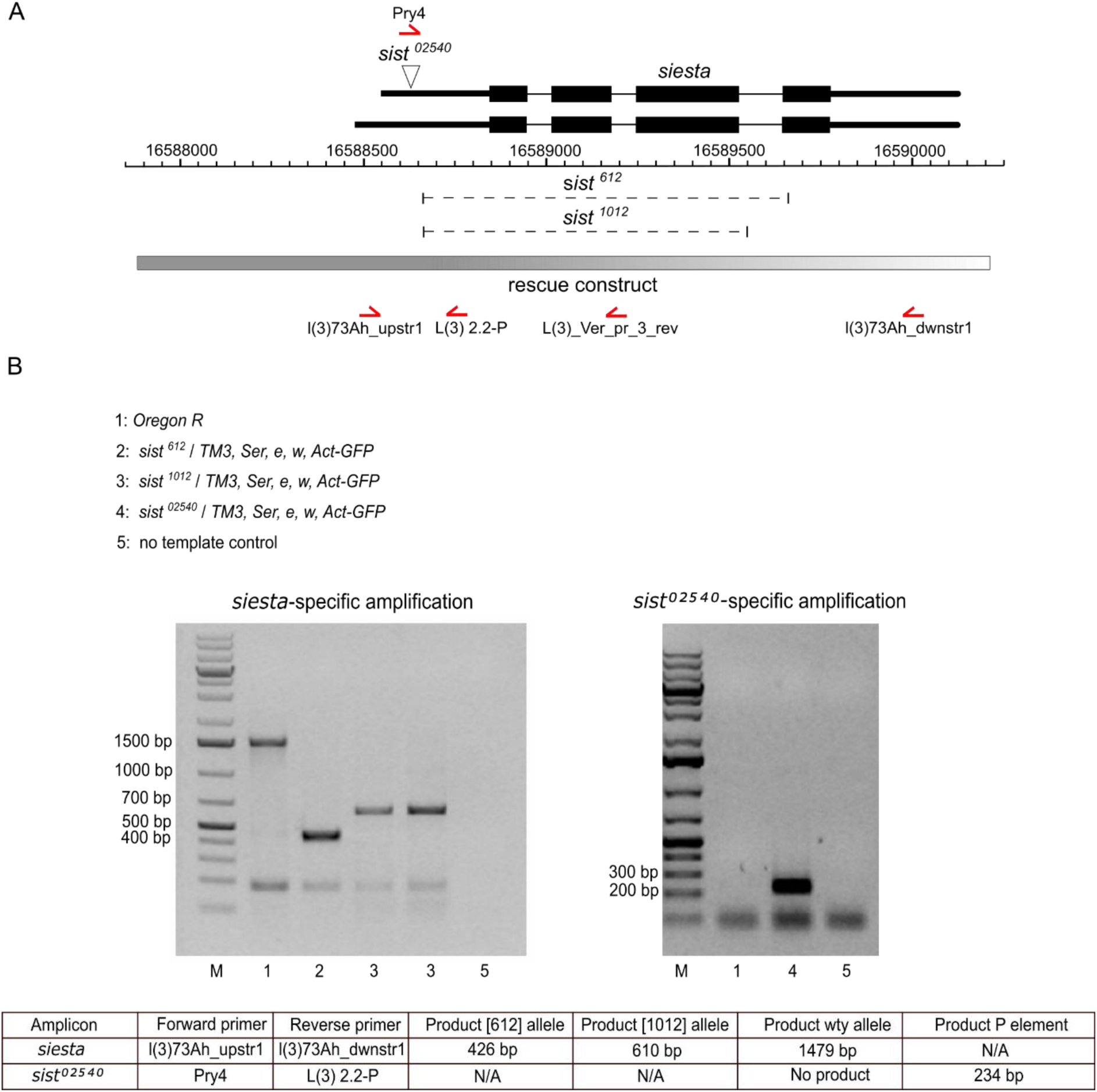
The structure and PCR genotyping of *Siesta* alleles. **A.** The schematics of the *siesta* locus. Two alternative transcripts above the coordinate scale (*dm6* genomic release) are shown with Transcription Start Sites (TSS) to the left. Thin lines indicate introns, and black boxes correspond to the coding parts. The position of transposon insertion in the *sist^02540^* allele (also known as *l(3)73Ah^02540^*) is indicated by a white triangle. Dashed lines mark the extent of *sist^612^*and *sist^1012^* deletions. The grey box indicates the genomic fragment sufficient to complement the *sist* loss-of-function mutations. Red half-arrows indicate positions of PCR primers used for genotyping. **B.** To verify the presence of specific *siesta* alleles, genomic DNA from flies of the indicated genotypes was used as a template for PCR with the primer combinations shown in the table. DNA from Oregon-R strain was used as a control. PCR products were analysed by gel electrophoresis in 1% agarose gel along with the Gene Ruler 1kb Plus molecular weight marker (M).

**Figure S2.**
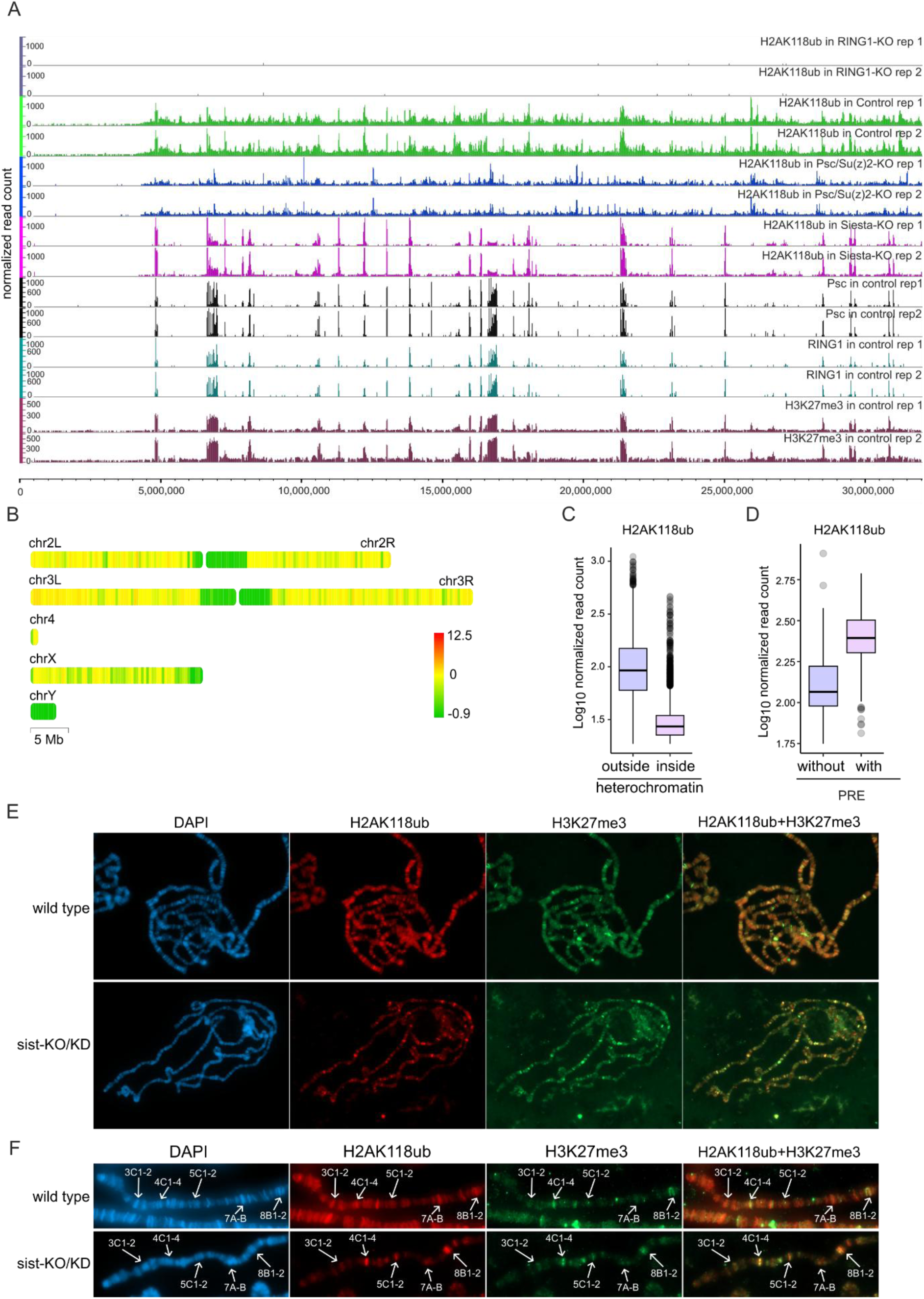
Overview of genomic H2AK118 ubiquitylation. **A.** Genome browser tracks showing H2AK118ub ChIP-seq profiles across *Drosophila* chromosome 3R in RING1-KO, control, Psc/Su(z)2-KO, and Siesta-KO cell lines (two replicates each). Additional tracks display ChIP-seq profiles for Psc, RING1, and H3K27me3 in control cells (two replicates each). The x-axis indicates genomic coordinates in *dm6* genomic release scale. **B.** The distribution of H2AK118ub ChIP-seq signal along *Drosophila* chromosomes. Average scaled signal intensities for fixed 350 kb windows tiling the genome were calculated using chromoMap v0.3 (Anand and Rodriguez Lopez, 2022). These were further converted into Z-scores and displayed as colour gradient ranging from green (low signal) to yellow (intermediate) and red (high signal). Note depletion of H2AK118ub from pericentromeric regions of chromosomes 2, 3 and X, as well as heterochromatin chromosome Y. Boxplots comparing normalized H2AK118ub ChIP-seq signals outside versus inside pericentromeric heterochromatin regions (**C**), and at genomic regions with or without Polycomb Response Elements (PREs) (**D**). Boxplots indicate the median and span interquartile range with whiskers extending 1.5 times the range and outliers shown as circles. **E.** Representative pictures of polytene chromosome spreads from wild type (*Oregon R*) and *sist^02540^*/*sist^612^* (Sist-KO/KD) third instar larvae stained with DAPI and antibodies against H2AK118ub and H3K27me3. The rightmost images show the merge between two immunostainings. **F.** Representative pictures of the distal part of X chromosome from the third instar larvae of the same genotypes. Positions of characteristic polytene chromosome bands according to Bridges nomenclature (Bridges, 1935) are marked with white arrows. Note the loss of H2AK118ub signal from the chromosome arms in the Sist-KO/KD mutants except from sites brightly stained with H3K27me3.

**Figure S3.**
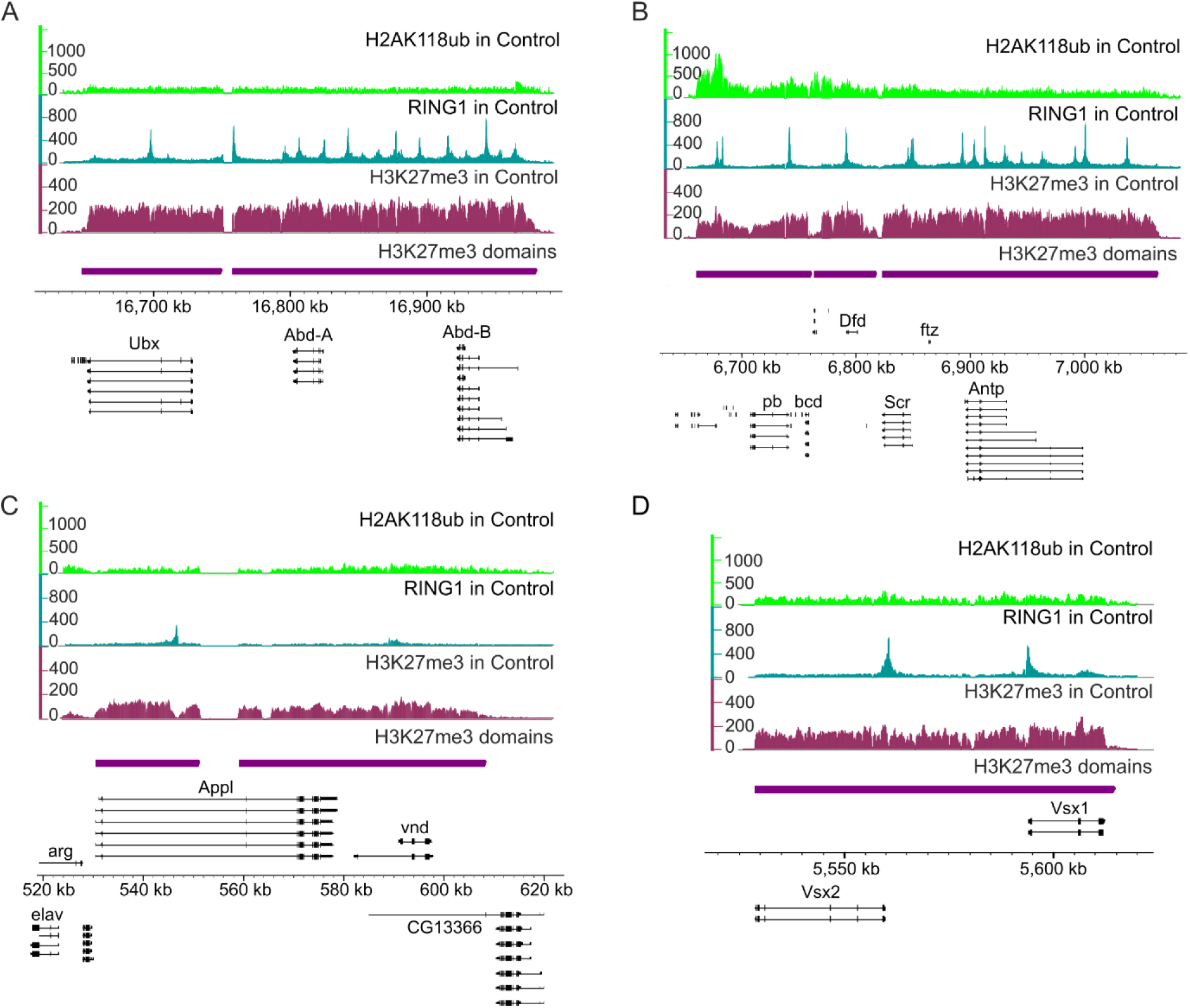
Examples of loci repressed by the Polycomb system with very low H2AK118ub. Genome browser tracks for H2AK118ub, RING1 and H3K27me3 ChIP-seq signals in control cells over the *bithorax complex* (**A**), the *Antennapedia complex* (**B**) *Appl – vnd* locus (**C**) and *Vsx1 – Vsx2* gene cluster (**D**). Genes shown above the coordinates scale (in *dm6* genomic release) are transcribed from left to right, genes shown below the coordinates scale are transcribed from right to left. The extent of H3K27me3 domains used to compare H3K27me3 and H2AK118ub ChIP-seq signals is shown as thick purple lines below the genome browser tracks.

**Figure S4.**
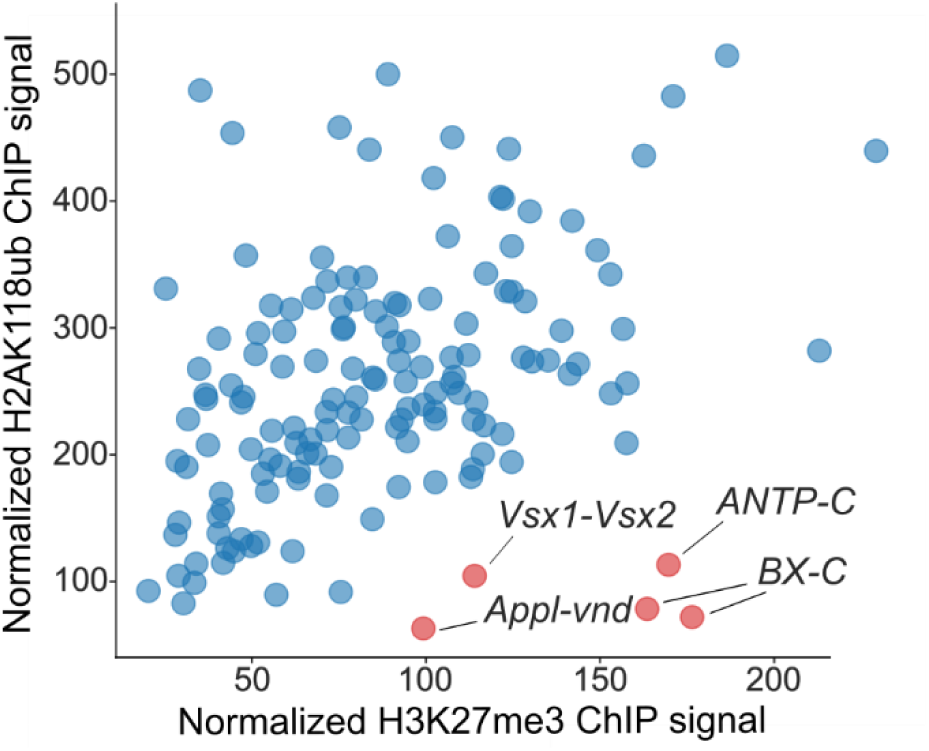
A small number of PRE-equipped genes have very low H2AK118ub compared to H3K27me3. The scatter plot shows averaged normalized ChIP-seq signals within continuous PRE-containing regions with significantly elevated H3K27me3 ChIP-seq signals (H3K27me3 domains). Each point corresponds to one H3K27me3 domain.

**Figure S5.**
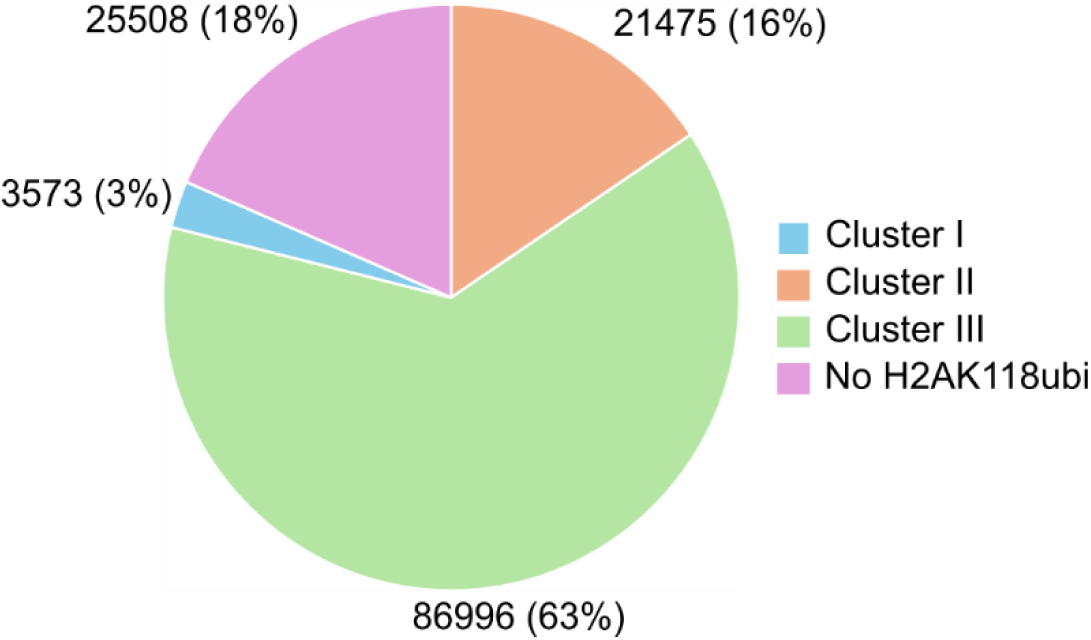
Relative representation of 1kb genomic segments (bins) assigned to four different categories based on comparison of H2AK118ub ChIP-seq signals in the control, Psc/Su(z)2-KO and Siesta-KO cells. An absolute number of 1kb bins in each category and percentile (in parentheses) is shown next to the corresponding segments of the pie chart.

**Figure S6.**
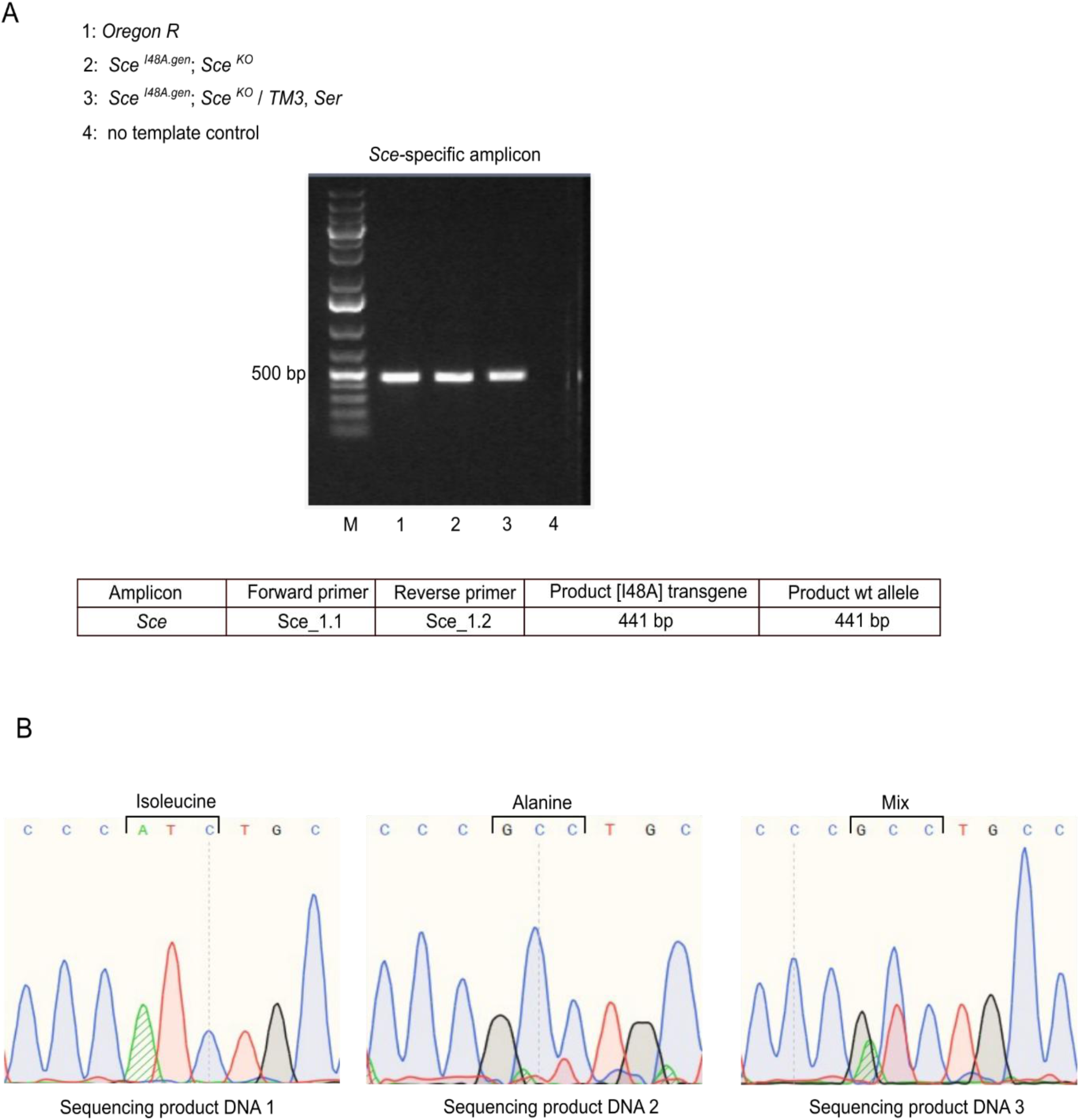
Genotyping of *Sce^I48A.gen^; Sce^KO^* (RING1-I48A) larvae. **A.** To verify the presence of *Sce* alleles, genomic DNA from flies of the indicated genotypes was used as a template for PCR using the primer combination shown in the table. DNA from Oregon-R strain was used as a control. PCR products were analysed by gel electrophoresis in 1% agarose gel along with the Gene Ruler 1kb Plus molecular weight marker (M). **B.** Chromatograms of sequencing reactions with PCR products from (**A**) show the Isoleucine to Alanine substitution at position 48 of the transgenic *Sce*.

**Figure S7.**
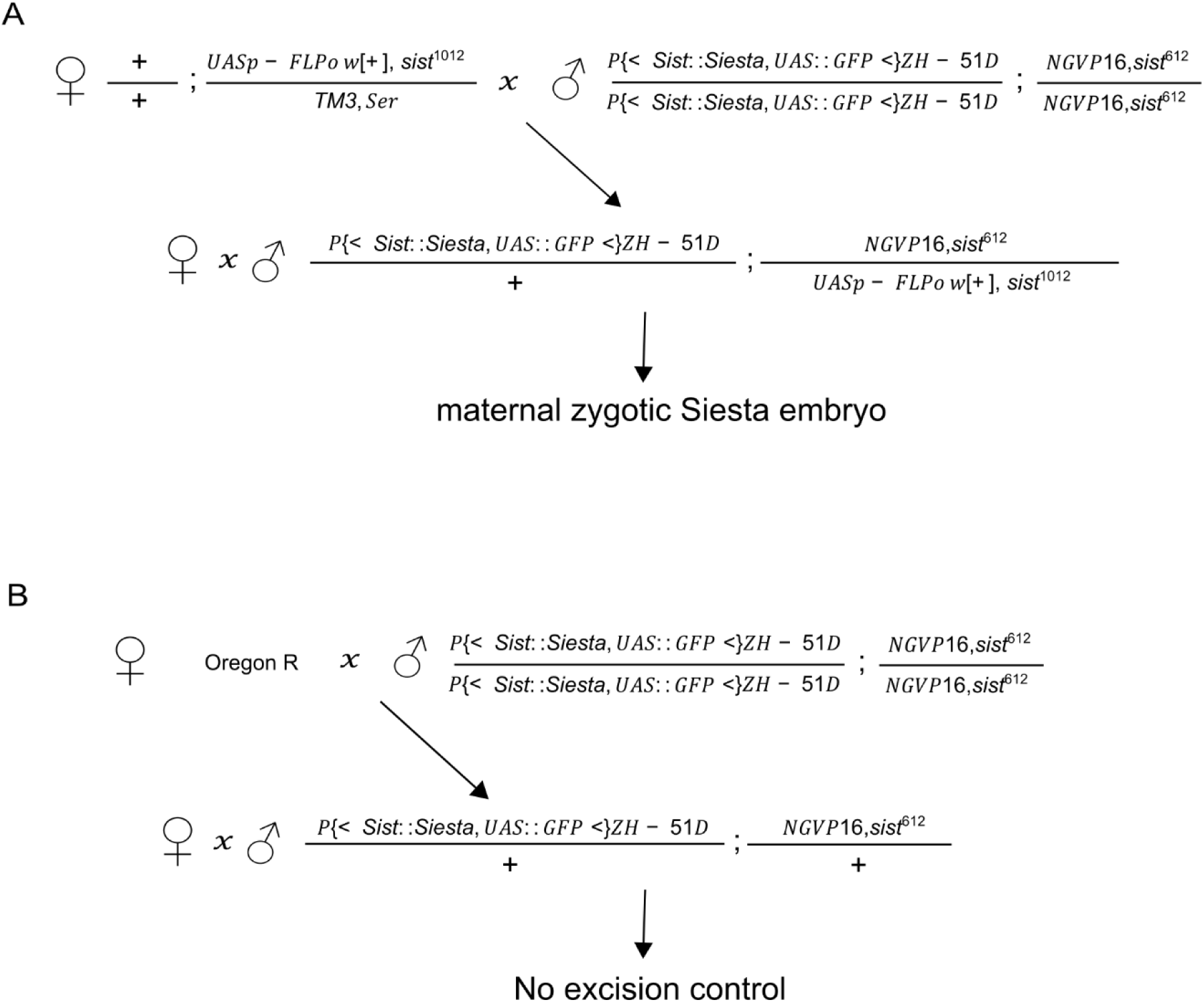
Generation of embryos that lack maternal and zygotic Siesta protein. **A.** Siesta KO^mz^ embryos were generated as described by (Gambetta and Muller, 2014). To this end, the *P{<sist::siesta, UAS::GFP<}ZH-51D; P{w^+mC^ = GAL4::VP16-nos.UTR}CG6325^MVD1^, sist^612^*males were crossed to the *w^−^/w^−^; +/+; sist^1012^, {UASp-FLPo, w^+^}VK33, y^+^/TM3, w, Ser, Act::GFP, e* females. From the resulting F1 progeny, males and females of the *P{<sist::siesta, UAS::GFP<}ZH-51D/+; P{w^+mC^ = GAL4::VP16-nos.UTR}CG6325^MVD1^, sist^612^/sist^1012^, {UASp-FLPo, w^+^}VK33, y^+^* genotype were crossed with each other, which resulted in embryos that lack maternal and zygotic Siesta protein. **B.** Control embryos were generated by crossing Oregon R females with *P{<sist::siesta, UAS::GFP<}ZH-51D; P{w^+mC^ = GAL4::VP16-nos.UTR}CG6325^MVD1^, sist^612^* males. The F1 progeny were then crossed with each other. Since there was no FLP recombinase source, the transgenic cassette was not excised.

**Figure S8.**
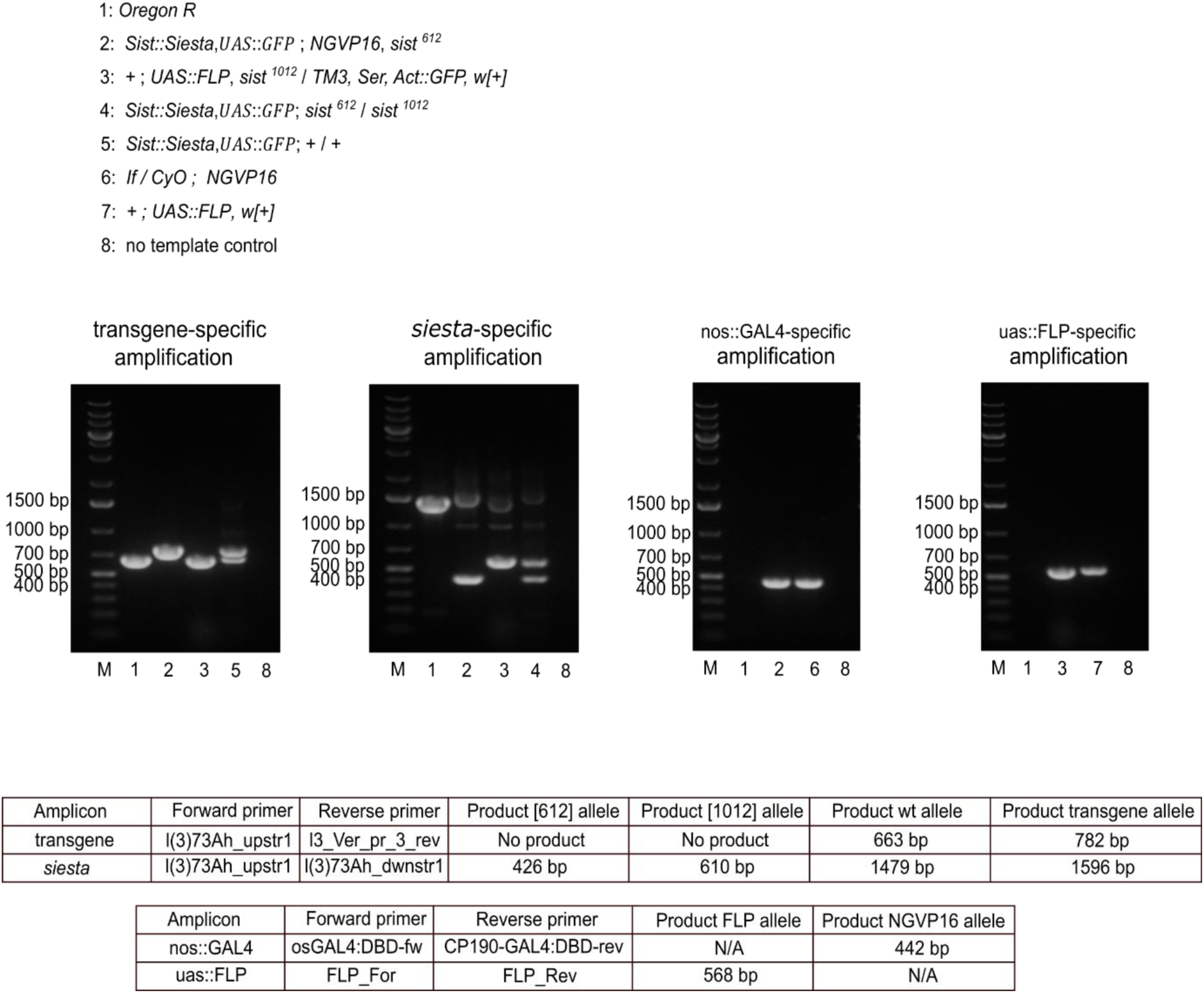
Genotyping of Drosophila strains involved in the generation of embryos lacking maternal and zygotic Siesta protein. DNA from flies of the indicated genotypes was used as a template for PCR using the primer combination shown in the tables. DNA from Oregon-R strain was used as a control. PCR products were analysed by gel electrophoresis in 1% agarose gel along with the Gene Ruler 1kb Plus molecular weight marker (M).

**Figure S9.**
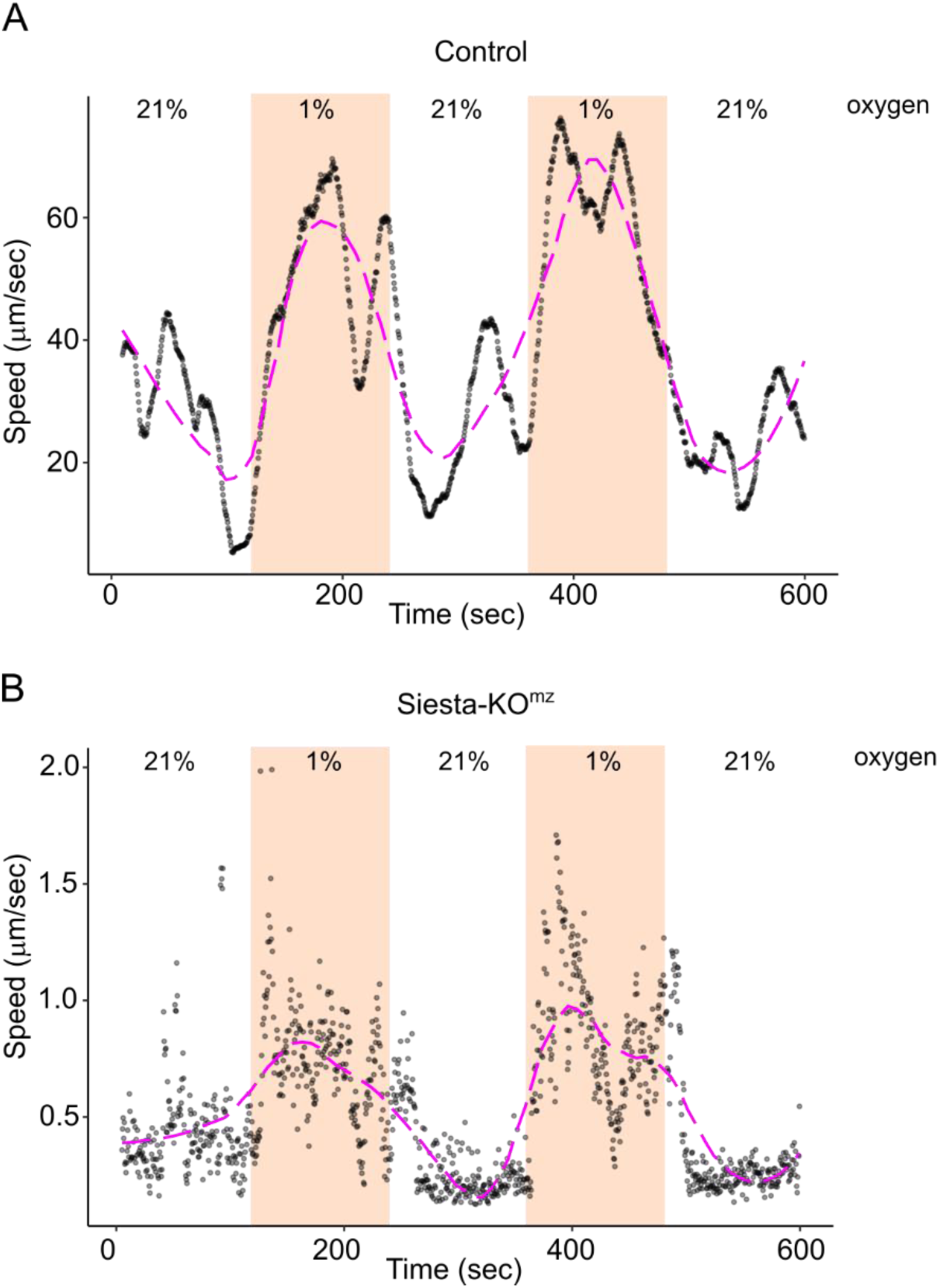
Siesta mutants respond to hypoxia. Motion tracking of control (**A**) and Siesta-KO^mz^ (**B**) first instar larvae under variable oxygen concentrations. The dots indicate the mean speed of all larvae in the camera view field at a given time point. The magenta dashed lines represent the outcome of data fitting with LOESS regression to demonstrate the general trend in speed changes. Note the different y-axis scales for the control and mutant larvae. Both control and Siesta-KO^mz^ larvae increase their crawling speed when exposed to 1% oxygen.

**Figure S10.**
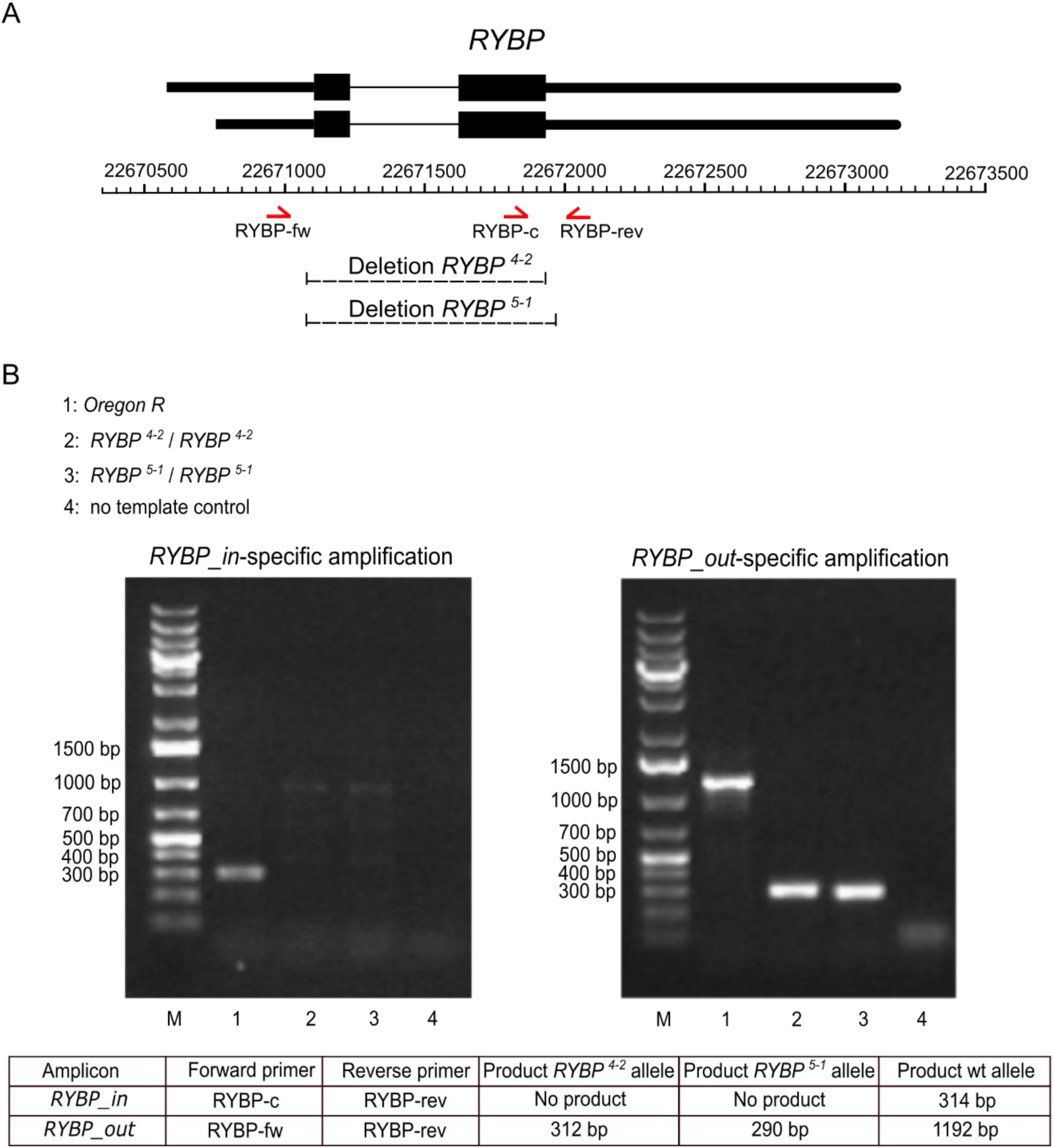
The structure and PCR genotyping of new *RYBP* alleles. **A.** The schematics of the *RYBP* locus. Two alternative transcripts above the coordinate scale (*dm6* genomic release) are shown with Transcription Start Sites (TSS) to the left. Thin lines indicate introns, and black boxes correspond to the coding parts. Dashed lines mark the extent of *RYBP^4-2^*and *RYBP^5-1^* deletions. Red half-arrows indicate positions of PCR primers used for genotyping. **B.** To verify the presence of specific *RYBP* alleles, genomic DNA from flies of the indicated genotypes was used as a template for PCR with the primer combinations shown in the table. DNA from Oregon-R strain was used as a control. PCR products were analysed by gel electrophoresis in 1% agarose gel along with the Gene Ruler 1kb Plus molecular weight marker (M).

**Figure S11.**
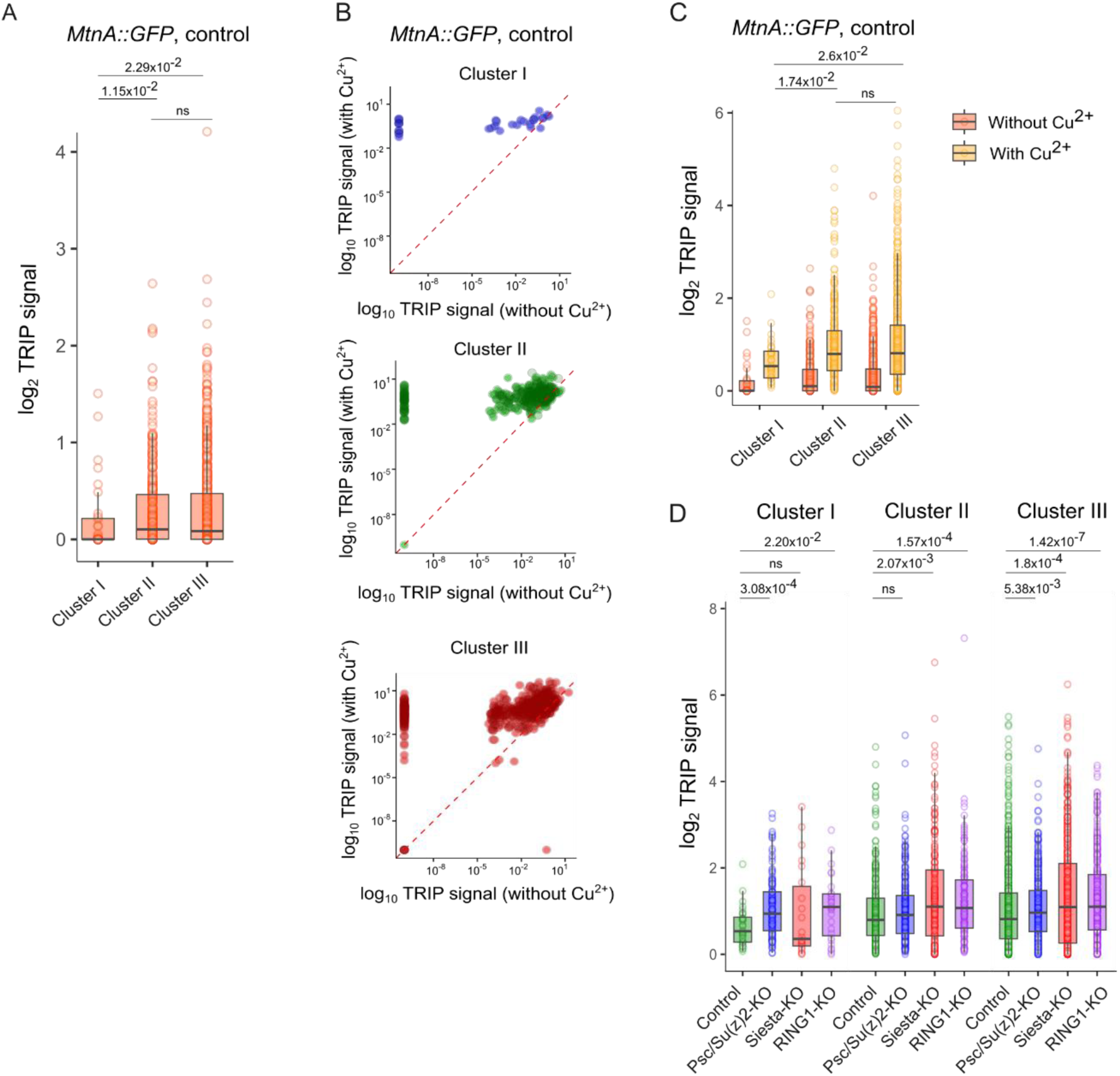
TRIP reveals no negative correlation between transcription and H2AK118ub. **A.** TRIP signals for the *MtnA::GFP* transgenes integrated in three types of genomic regions in control cells. Here and in **C** and **D**, the boxplots indicate the median and span interquartile range with whiskers extending 1.5 times the range and outliers shown as circles. The differences in medians between corresponding groups were tested for statistical significance using the Wilcoxon rank sum test, and p-values are displayed above the box plots. **B.** Log-log comparison of TRIP signals for the *MtnA::GFP* transgenes before and after copper Cu^2+^ induction in three types of genomic regions in control cells. A dashed red diagonal line (slope = 1) denotes equal signal before and after induction. **C.** TRIP signals for the *MtnA::GFP* transgenes integrated into three types of genomic regions in control cells before and after copper Cu^2+^ induction. **D.** TRIP signals for the *MtnA::GFP* transgenes in different cell lines.

**Figure S12.**
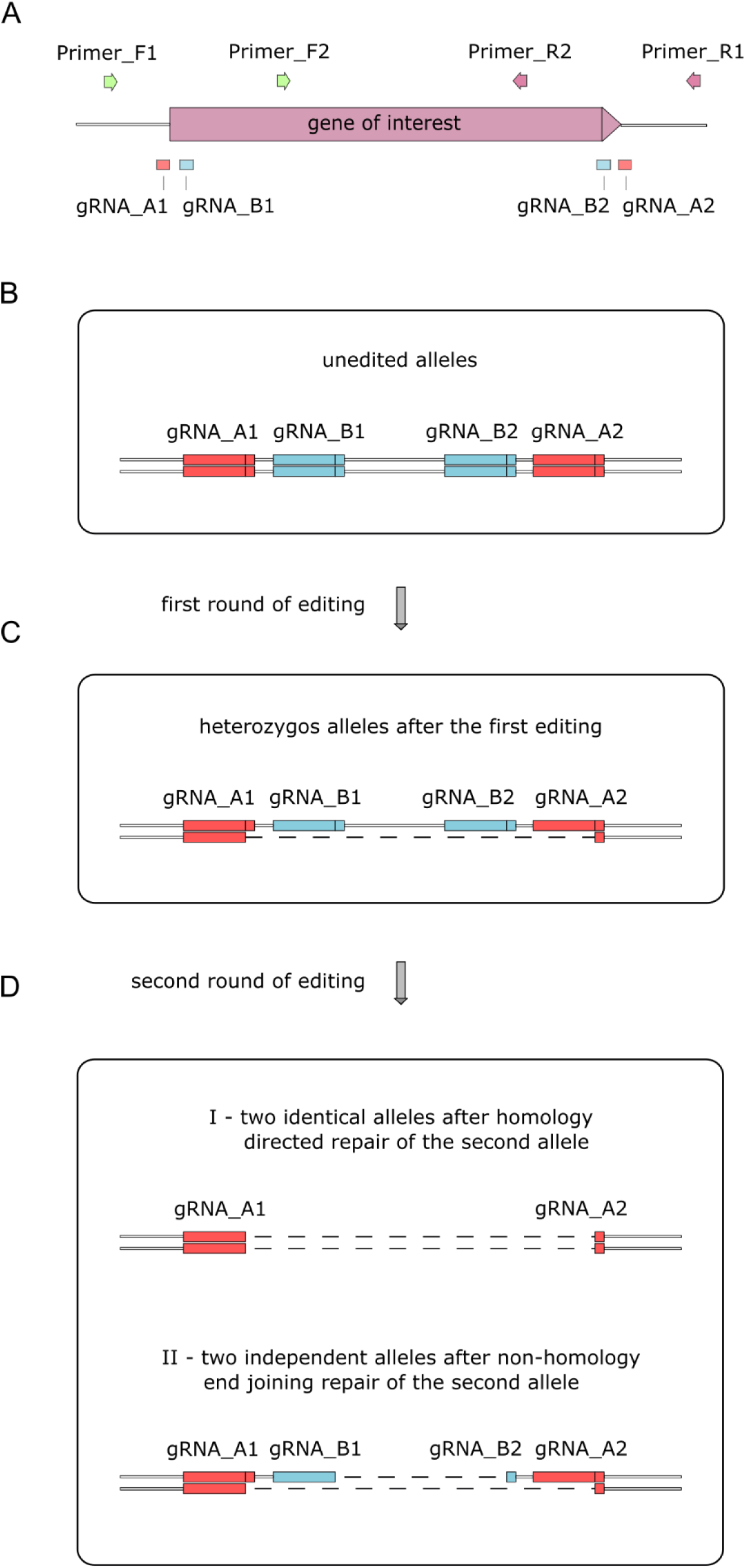
CRISPR/Cas9-mediated gene editing in cultured *Drosophila* cells. **A.** Schematics of a gene to be edited with locations of target sequences for gRNAs and primers for genotyping. gRNA_A1 and gRNA_A2 are used for the first round of editing, gRNA_B1 and gRNA_B2 are used for the second round of editing. Primer_F1 and Primer_R1 are used for amplification of the junction after DNA repair and sequencing. Primer_F2 and Primer_R2 are used to screen for the clones with homozygous deletion; no PCR product is expected in homozygous cells. **B.** The schematic of unedited alleles. Rectangles correspond to the position of target sequences for gRNAs, vertical lines inside each rectangle indicate the position of the Cas9 cut in case of the precise deletion. **C.** Combination of alleles in heterozygous cells after the first round of editing. Most of the selected single cell clones bear only one allele with a deletion. **D.** The two most frequent combinations of alleles in the resulting clones after the second round of editing.

**Figure S13.**
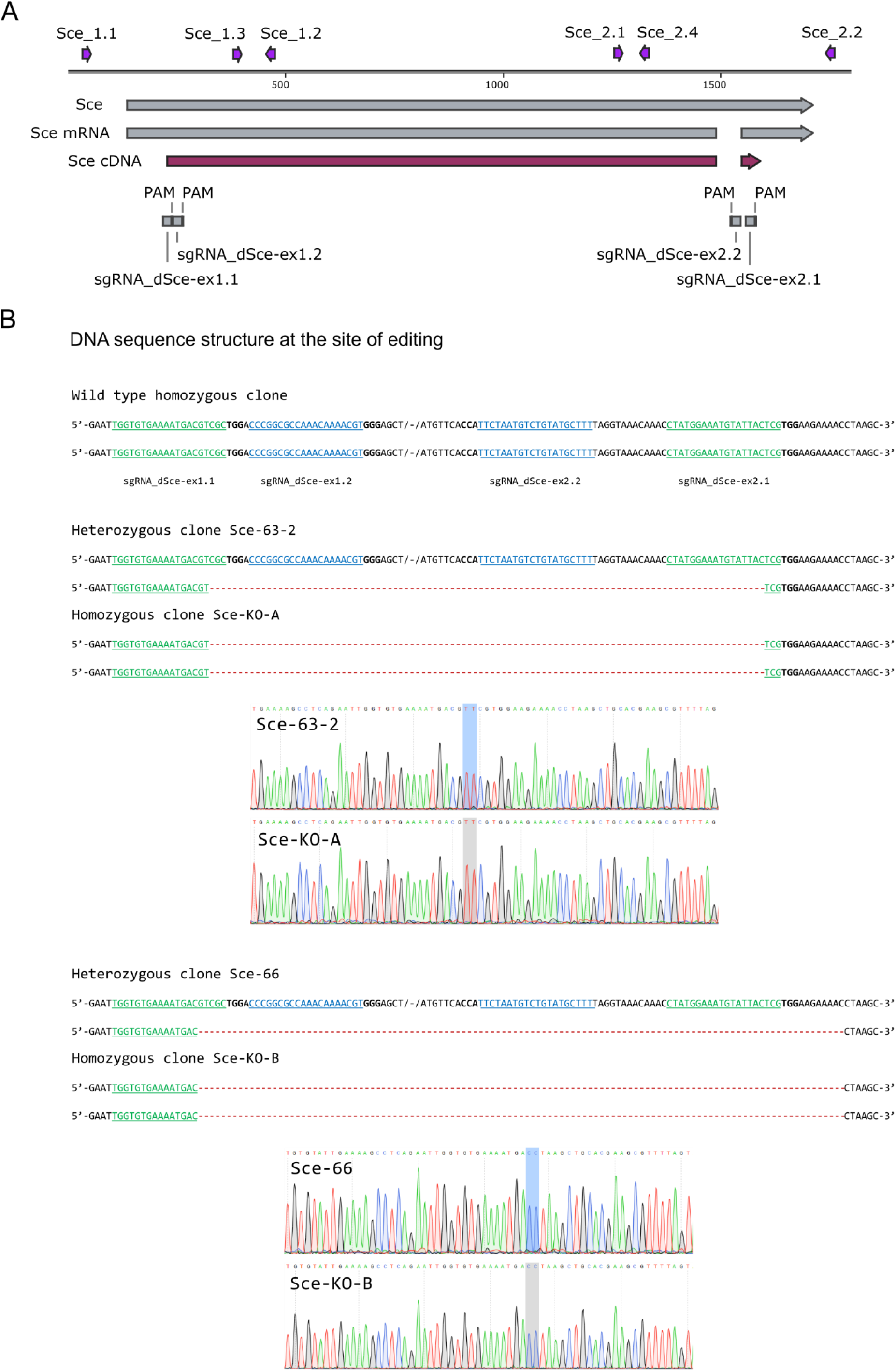
Generation of cultured RING1-KO Drosophila cells. **A.** The *Sce* gene structure and location of target nucleotide sequences for guide RNAs and primers. The *Sce* gene, mRNA and cDNA are represented as long arrows. Primer locations are indicated as pink arrows, locations of target nucleotide sequences for guide RNAs with corresponding Protospacer Adjacent Motifs (PAMs) are shown as grey boxes. The nucleotide sequences of PCR primers are listed in Table S3. **B.** DNA sequences of both homologous chromosomes at the editing site in unedited cells, heterozygous cells and homozygous cells from the resulting cell lines. The gRNA target sequences are underlined and shown in green for the outer pair of gRNAs, in blue for the inner pair of gRNAs. Triplets in bold correspond to PAMs. Red dashed lines mark the position of deleted nucleotides. Blue and grey vertical shadows on sequencing chromatograms mark the position of the junction.

