## Supplementary material for "Polycomb repression works without Siesta": Supplementary_file_7.txt

# awk based bash script for demultiplexing of TRIP data, replace extension of the file name with .sh to execute
# Polycomb repression works without Siesta; Tatyana G. Kahn, Andres Garrido, Anastasiya Yushkova, Maria Kim, Alexander Glotov, Sweda Sreekumar, Jan Larsson, and Yuri B Schwartz

#!/bin/bash

# dataset of FASTQ files to process
files=(
    "Ras17-N2_1.fastq"
    "Psc-N1_1.fastq"
    "Psc-N2_1.fastq"
    "Sce-N1_1.fastq"
    "Sce-N2_1.fastq"
    "L10-N1_1.fastq"
    "L10-N2_1.fastq"
    "Ras17-N1_1.fastq"
)

# function to process FASTQ files with specific promoter index inclusion and exclusion rules
process_fastq() {
    local pattern_include="$1"  # promoter index pattern to include in sequences
    local suffix="$2"           # suffix for output filenames based on the barcode

    # define exclusion pattern based on the selected index
    if [[ "$pattern_include" == "ACTAG" ]]; then
        pattern_exclude="ACCAT|TACGT|TTGTA"
    elif [[ "$pattern_include" == "ACCAT" ]]; then
        pattern_exclude="ACTAG|TACGT|TTGTA"
    elif [[ "$pattern_include" == "TACGT" ]]; then
        pattern_exclude="ACTAG|ACCAT|TTGTA"
    elif [[ "$pattern_include" == "TTGTA" ]]; then
        pattern_exclude="ACTAG|ACCAT|TACGT"
    else
        echo "Invalid pattern: $pattern_include"
        return
    fi

    for file in "${files[@]}"; do
        output_file="${file%.fastq}_$suffix.fastq" # create output filename

        awk -v pat_include="$pattern_include" -v pat_exclude="$pattern_exclude" '
            NR % 4 == 1 {header = $0; next}         # store the header line
            NR % 4 == 2 {                           # sequence line
                seq = substr($0, 50, 21);            # extract from 50 to 70 bp
                if (seq ~ pat_include && seq !~ pat_exclude) {  # match inclusion & exclude unwanted
                    print header;                   # print identifier line
                    print $0;                       # print full sequence
                    getline; print $0;              # print "+" separator
                    getline; print $0;              # print quality line
                }
            }
        ' "$file" > "$output_file"
    done
}

# process each promoter index and assign a suffix
process_fastq "ACTAG" "11"
process_fastq "ACCAT" "14"
process_fastq "TACGT" "3"
process_fastq "TTGTA" "16"

echo "Processing complete."
