## Supplementary material for "Polycomb repression works without Siesta": Supplementary_file_8.txt

# Configuration file for TRIP analysis based on TRIP Analysis Software Kit (TASK) software (Akhtar, W., de Jong, J., Pindyurin, A.V., Pagie, L., Meuleman, W., de Ridder, J., Berns, A., Wessels, L.F., van Lohuizen, M., and van Steensel, B. (2013))


# Polycomb repression works without Siesta. Tatyana G. Kahn, Andres Garrido, Anastasiya Yushkova, Maria Kim, Alexander Glotov, Sweda Sreekumar, Jan Larsson, and Yuri B Schwartz


# Illumina sequencing was performed as paired-end, for the "expression" and "normalization" datasets, only single-end reverse 150bp reads containing a constant plasmid region and a unique barcode were used
# The third dataset contains "mapping" reads. These are paired-end 150bp reads obtained by sequencing of inverse PCR products. 
# The “mapping” reads include the 5’ end of the piggyBac transposon and a neighbouring genomic DNA sequence in the forward read, as well as a barcode and a genomic sequence adjacent to the DpnII site in the reverse read.

					
# the length of index seqeunce. If no index is part of the read (formats Exp2, Exp3, map2 and map3), then right 0.
# please note that the read structure at the start
 
index_length=	0	


# the length of the barcode. The program looks for barcodes which are barcode_length ± 1. 

barcode_length= 18


# the first constant part (pat1) in the normalization/expression reverse read. If the read is starting. 
# directly from barcode then rigth NA to this pattern. 
# Only ACTG are allowed.

pat1=	CGCCAGGGTTTTCCCAGTCACAAGGGCCGGCCACAACTC


# the second constant part (pat2) in the normalization/expression reverse read. Please note that XXXXX corresponds to a specific promoter index:
# "XXXXX" = "ACTAG" = index "11", pPB-Hsp70-eGFP-PI-11-BC-libr 
# "XXXXX" = "ACCAT" = index "14", pPB-MtnA-eGFP-PI-14-BC-libr
# "XXXXX" = "TACGT" = index "3", pPB-Hsp70-eGFP-PI-3-BC-libr 
# "XXXXX" = "TTGTA" = index "16", pPB-MtnA-eGFP-PI-16-BC-libr

pat2=	CCTCXXXXXCTCGATC


# The maximum Hamming distance to filter out potential mutants. An hd of 2 will mean that 
# for a frequent barcode all other (less frequent) barcodes only 2 Hamming distance apart
# will be discarded from the list of genuine barcodes.

hd=	0


# the first constant part (map_pat1) in the reverse mapping read. If the read is starting. 
# directly from barcode then rigth NA to this pattern.
# Only ACTG are allowed.

map_pat1=	GTCACAAGGGCTGGCCACAACTC


# the second constant part in the reverse mapping read (map_pat2). Please note that XXXXX corresponds to a specific promoter index:
# "XXXXX" = "ACTAG" = index "11", pPB-Hsp70-eGFP-PI-11-BC-libr 
# "XXXXX" = "ACCAT" = index "14", pPB-MtnA-eGFP-PI-14-BC-libr
# "XXXXX" = "TACGT" = index "3", pPB-Hsp70-eGFP-PI-3-BC-libr 
# "XXXXX" = "TTGTA" = index "16", pPB-MtnA-eGFP-PI-16-BC-libr

map_pat2=	CCTCXXXXXCTCGATC


# the constant part in the forward mapping read. Only ACTG are allowed.

map_pat_rev=	GTACGTCACAATATGATTATCTTTCTAGGGTTAA


# the number of processors to be used for Bowtie2 alignments 

cores=	8


# the index of the genome to align the mapping reads against

bowtie_base=	/Dm6


# the maximum distance to cluster the positions on the forward read. Typically it is kept 
# a few hundered bases because during inversePCR of TRIP, two different closely spaced
# restriction sites can be used to make the product.

max_dist_for=	                500


# the maximum distance to cluster the positions on the reverse read. Typically it is kept 
# a few bases just to combine those reads for which the aligment was slightly shifted 
# because of sequencing errors.

max_dist_rev=	20


# the minimum number of reads for considering a barcode genuine. An arbitrary recommendation 
# is 5 reads. But if your sequencing depth is low than you might want to reduce it to 3 or 4.

min_counts= 2
