## Supplementary material for "Polycomb repression works without Siesta": Supplementary_file_9.txt

# To run the TASK software on a single dataset, use the following command in the terminal. In this example, the control dataset with index 11 is used:

perl trip0.5.pl -n ./Ras17-N1_1_11.fastq -e ./R17-T1_1.fastq -f ./Ras17-M1_1.fastq -r ./Ras17-M1_2.fastq -c config11.txt -o output -m r -v -d --out_dir ./index11/Ras_M1_N1_T1
